# Climate adaptation in *P. trichocarpa*: key adaptive loci identified for stomata and leaf traits

**DOI:** 10.1101/2024.07.11.603099

**Authors:** Marie C Klein, Zi Meng, Jack Bailey-Bale, Suzanne Milner, Peicai Shi, Wellington Muchero, Jin-Gui Chen, Timothy J Tschaplinski, Daniel Jacobson, John Lagergren, Matthew Lane, Chris O’Brien, Hari Chhetri, Mengjun Shu, Peter Freer-Smith, Thomas N. Buckley, Troy Magney, J Grey Monroe, Gerald A. Tuskan, Gail Taylor

## Abstract

Identifying the genetic basis of traits underlying climate adaptation remains a key goal for predicting species responses to climate change, enabling the elucidation of gene targets for future climate-resilient crops. Here, we measured 14 leaf and stomatal traits under control (well-watered) and drought conditions, subsampling a diversity collection of over 1,300 *Populus trichocarpa* genotypes, a potential biofuel feedstock crop. Stomatal traits were correlated with the climate of origin for genotypes, such that those originating from environments subject to water deficit tended to have smaller stomata, but with higher density. Stomatal traits were also correlated with leaf morphology, with larger leaves having larger stomata and lower stomatal density mirrored in correlations to climate of origin. The direction of plastic responses - reduced stomatal size under drought - mirrors the correlations seen among genotypes with respect to the aridity of environmental origin. Genome-Wide Association Studies (GWAS) identified loci underlying trait diversity, including candidates contributing to stomatal size. We used climate of origin to predict stomatal size in genotypes with unknown trait values and found that these predicted phenotypes confirmed empirically measured allele effects. Finally, we found evidence that future climates may select for alleles contributing to decreased stomatal size, with the strength of selection depending on the availability of moisture. These findings reveal adaptive variation in stomatal and physiological traits along with underlying genetic loci, with implications for future selection and breeding - providing insights into the responses to future climate change.

**Highlight:** Research on *Populus trichocarpa* reveals adaptation of physiological and stomatal traits linked to drought tolerance, with genotypes from arid regions exhibiting smaller stomata, offering insights for climate change adaptation and sustainable biofuel production.

## Introduction

Altered rainfall patterns and rising temperatures are intensifying droughts, jeopardizing agricultural output due to scarce irrigation water (Cook et al., 2018; Dai, 2012; Food and Agriculture Organization of the United Nations et al., 2018). Moreover, in anticipation of the emerging circular bioeconomy, there is a growing need for fast-growing non-food trees and grasses (Clifton-Brown et al., 2019; Somerville et al., 2010) that can both grow on marginal lands (Hoegh-Guldberg et al., 2018; Mehmood et al., 2017; Schmidt et al., 2015) and tolerate drought and limited nutrient inputs. Consequently, a pivotal challenge is to achieve economically viable yields on marginal lands, in the face of limited water availability (Hoegh-Guldberg et al., 2018; Taylor et al., 2019). Given these challenges, it is crucial to study the physiology and genetics of climate adaptation in potential biofuel crops, like poplar (*Populus* spp). Understanding the genetic basis and adaptive value of ecophysiological traits can facilitate the development of varieties that are resilient to arid climates (Blumstein et al., 2020; Savolainen et al., 2013; Stapley et al., 2010).

Collections of diverse genotypes of natural populations, originating from wide-ranging environments, are invaluable resources to address these challenges (Taylor et al., 2024). Such collections enable the identification of links between traits and their climates of origin, shedding light on the adaptive value of phenotypic diversity. This can inform predictions of which traits are suitable for dry climates and guide the selection of genotypes for cultivation in arid regions. Additionally, the genetic basis of these traits can be elucidated through this genomic diversity. This exploration is essential for pinpointing genetic markers that can be used to accelerate breeding programs and conservation efforts. Stomatal traits are central to plant water management and exhibit natural variation in features including size and density, which have been linked to climate of origin (Beerling & Woodward, 2008; Franks & Beerling, 2009). Smaller and denser stomata, for instance, have often been observed in genotypes from drier climates, suggesting an evolutionary adaptation to aid water conservation and improve plant water-use efficiency (WUE) (McKown et al., 2019; McKown, Guy, Klápště, et al., 2014; McKown, Guy, Quamme, et al., 2014). This link underscores the importance of stomata in local adaptation and WUE strategies (Dittberner et al., 2018; Doheny-Adams et al., 2012; Franks et al., 2009; Kardiman & Ræbild, 2018; Ohsumi et al., 2007; Sun et al., 2014). However, the scope of leaf traits influencing water dynamics extends beyond stomata (McKown, Klápště, et al., 2014). Leaf area, leaf mass, specific leaf area (SLA), photosynthetic potential, surface wettability, and integrated rather than instantaneous WUE form an intricate network that characterizes plant interaction with the environment. Each trait contributes to a plant’s ability to conserve water, optimize photosynthesis, and survive in varying conditions of water availability (Benavides et al., 2021). For example, smaller leaves with higher SLA may transpire less water and larger leaves might capture more light, balancing the trade-offs between water conservation and energy acquisition (Franks & Beerling, 2009; Hetherington & Woodward, 2003; Z. Liu et al., 2020; Wright et al., 2004, 2017; Wu et al., 2016). Similarly, wettability can affect leaf temperature and hence water loss through transpiration, while variations in photosynthetic rates can reflect different strategies to maximize carbon gain against water loss (Nobel, 1999). At the same time, the direct correlation between total plant water use and yield is also well-established, including for poplar, with high-yielding ‘water spending’ and lower-yielding ‘water saving’ strategies varying in their usefulness depending on the intensity, longevity, and life-stage of the drought (Tardieu, 2022). Unraveling these conflicting ecological strategies remains challenging.

To assess the degree of local adaptation of leaf and stomatal traits, we established a common garden of black cottonwood (*Populus trichocarpa*) in Davis, California, an extreme hot and dry site. These trees are derived from a natural population from the Pacific Northwest, representing a wide spectrum of climates (Evans et al., 2014; Gornall & Guy, 2007; McKown, Guy, Klápště, et al., 2014). Recognized for its fast growth and high cellulose content, *Populus trichocarpa* stands out as a promising bioenergy crop, including for liquid biofuel production alongside Bioenergy with Carbon Capture and Storage (BECCS). In addition, *Populus* is a valuable model for genetics and plant biology research (Kačík et al., 2012; Taylor et al., 2019; Tuskan et al., 2006). Such attributes augment the significance of *P. trichocarpa* trees in achieving the U.S. Department of Energy’s ambitious Sustainable Aviation Fuel (SAF) target by 2050, while simultaneously reducing life cycle greenhouse gas emissions by at least fifty percent (U.S. Department of Energy, 2022).

Although *Populus* trees, as riparian plants, are typically considered vulnerable to drought, notable differences in drought response have been documented among *Populus* genotypes (Cocozza et al., 2010; Huang et al., 2009; Marron et al., 2002; Monclus et al., 2006; Regier et al., 2009; Street et al., 2006; Tschaplinski et al., 1994, 2006; Viger et al., 2013) and the genus can be found across contrasting climate zones, including in extremely arid environments (Brosché et al., 2005). Further research is needed to elucidate the physiological and genetic basis of drought tolerance in these trees to enable tree establishment and productivity in poor and marginal environments (Taylor et al., 2019). In this study, we investigated the role and importance of 14 leaf physiology traits related to leaf morphology, stomata, and water use in *P. trichocarpa*, under both drought and well-watered conditions. twice during the growing season in a subsample of approximately 3,000 trees of 469 unique genotypes (Figure 1, Table 1) and measured a biomass yield trait in the whole population (approximately 750o tress, 1382 unique genotypes. In doing so, we explore the adaptive signature of drought tolerance and plastic responses in this widely divergent population. To overcome the difficulties associated with phenotyping a large number of samples, we employed a suite of automated phenotyping algorithms for data acquisition. Among leaf traits, we also assessed δ^13^C — a metric for WUE in wood sampled during tree dormancy, shown in *Populus* to be valuable and correlated with WUE (Bogeat-Triboulot et al., 2019; Viger et al., 2016). The goal of our study was to explore the relationship between trait variation and climate of origin, assessing the adaptive value of trait variation, their relevance and usefulness to trait selection for drought tolerance, and the roles of plastic, acclimatory responses. Finally, genome-wide association analyses were conducted to characterize the genetic basis of these traits, enabling a more holistic view of climate-genotype-trait relationships in *P. trichocarpa* that reflects local adaptation and provides gene targets for future tree improvement for the emerging bioeconomy.

**Figure 1.**
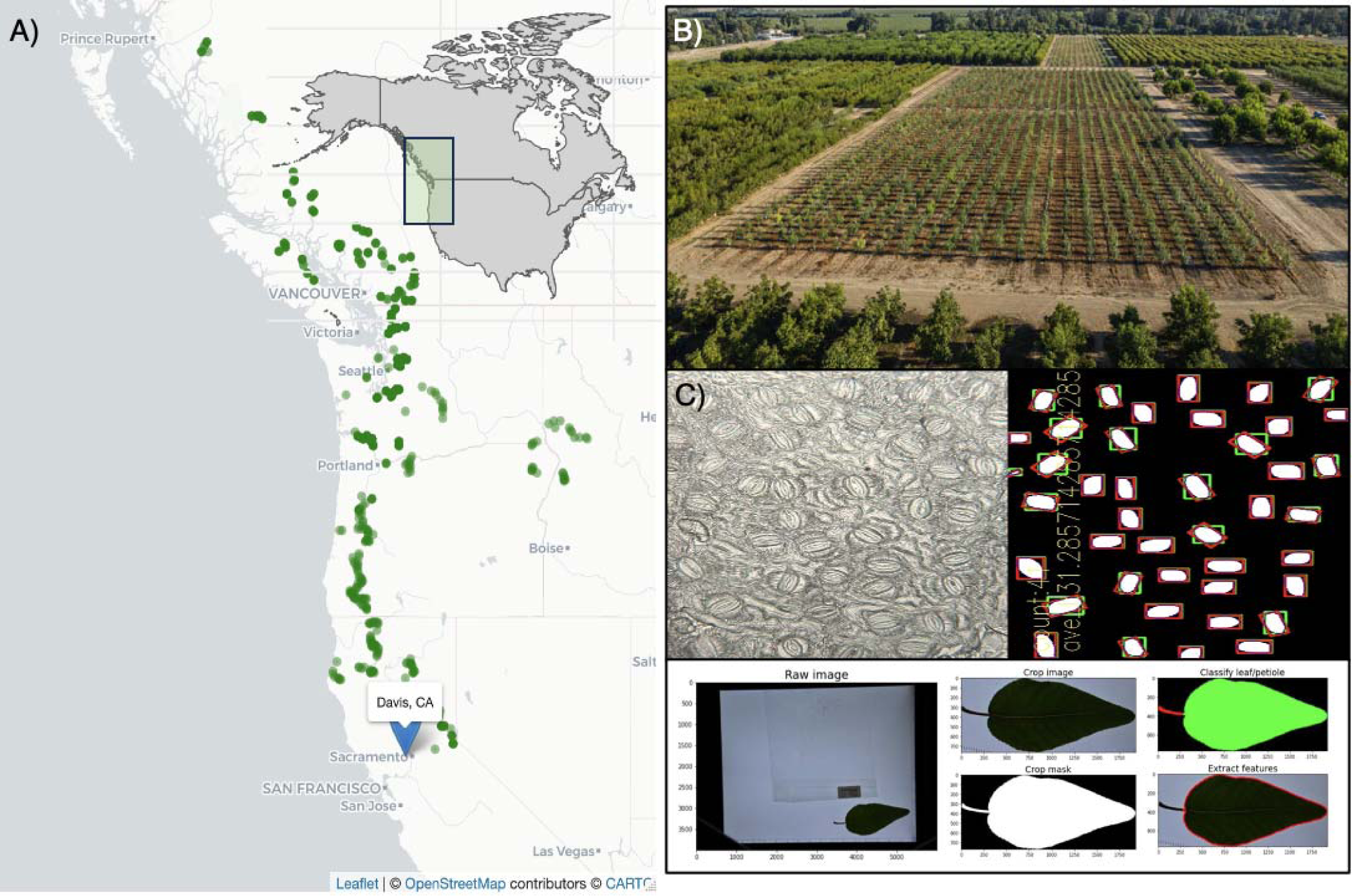
Overview of experiment. A) Location of *P. trichocarpa* collections in the Pacific North B) Common garden located in Davis, CA., and C) Examples of Stomatal (above) and leaf phenot (below): Stomata phenotypes: On the left: Raw microscopy image; on the right: Processed image suitable for identifying stomatal density and size, captured at 16x magnification. Leaf phenotypes: Examples illustrating the extraction of leaf area and perimeter measurements from raw image (left).

**Table 1.**
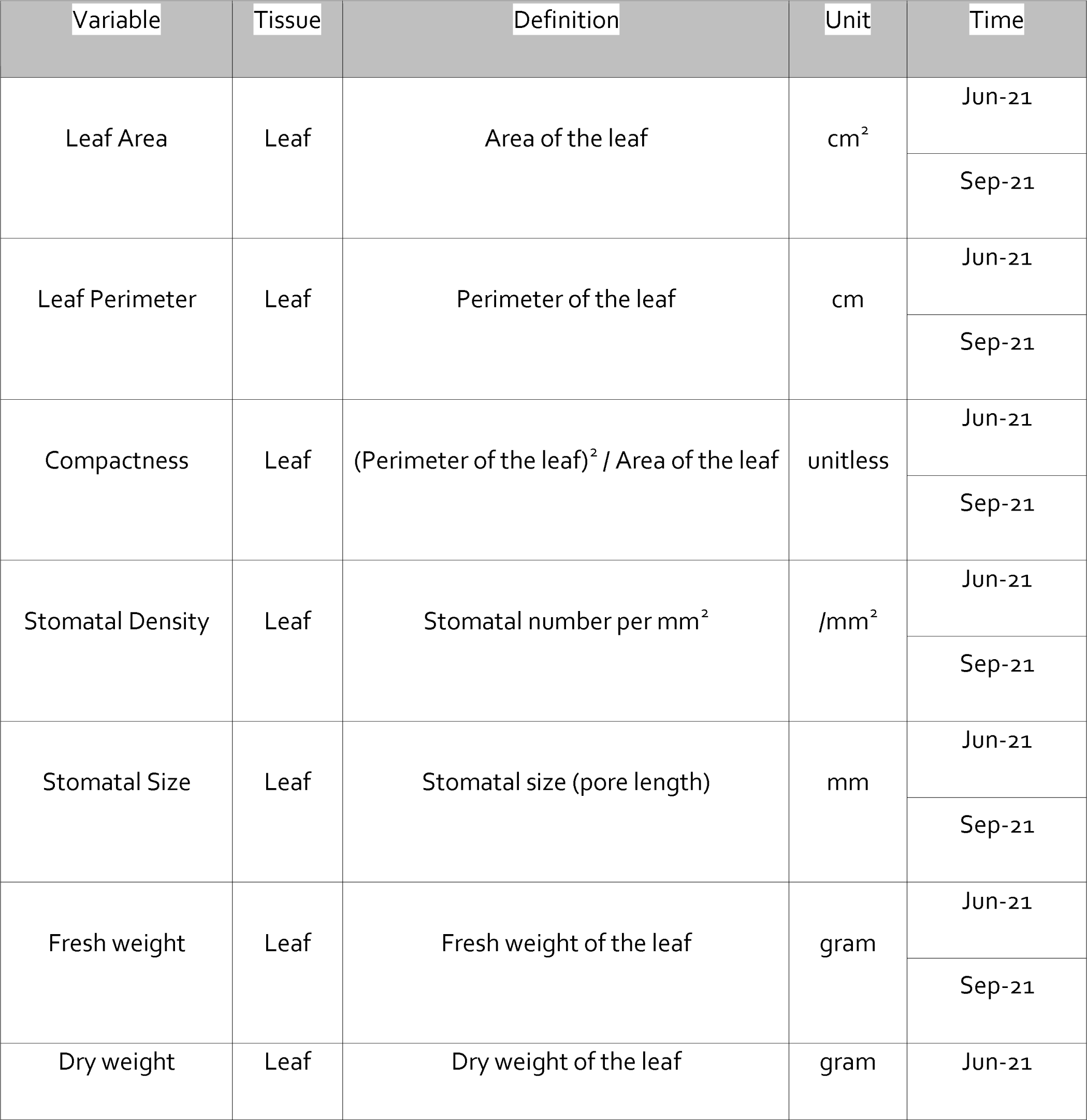
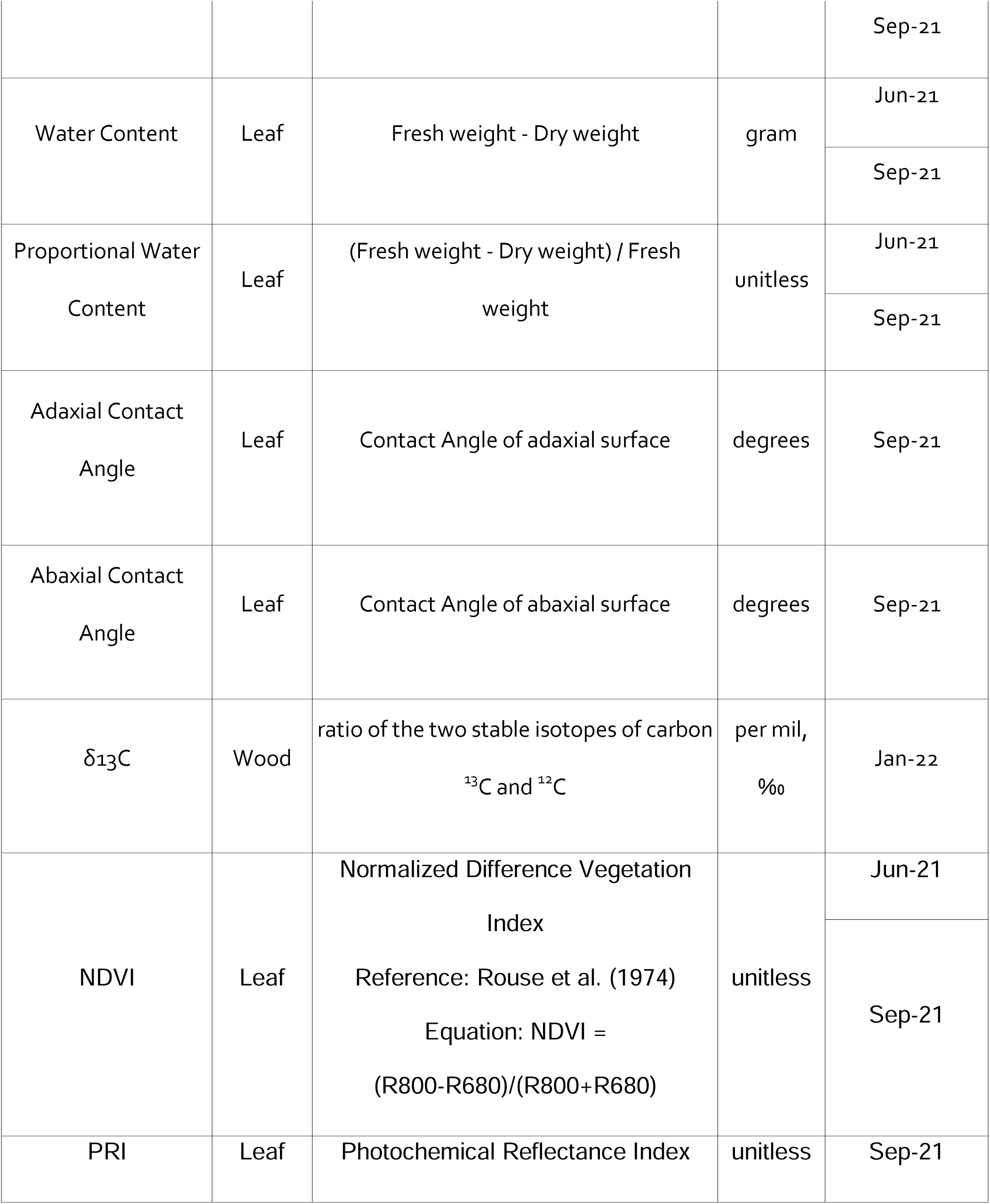
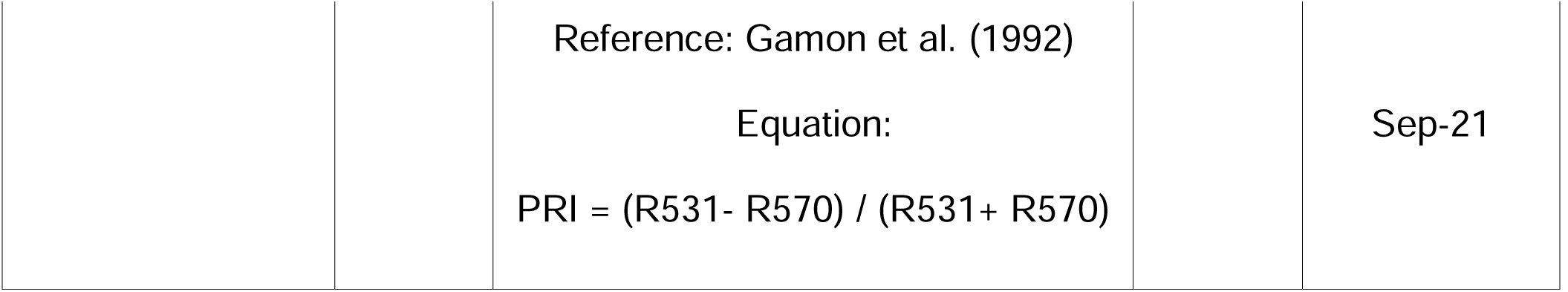
Phenotypes measured and used for analysis.

## Materials and Methods

### Field Site and Plant Material

The research was conducted on a 6.1-hectare plot at the UC Davis Plant Science Field Facility in Davis, CA (38°32’47.4’’N, 121°47’32.7’’W) (**Figure 1B**) as described previously (Taylor et al., 2024). The experimental layout consisted of drought and control treatments across adjacent fields that were divided into three distinct blocks, arranged in an incomplete randomized block design. The *P. trichocarpa* population used in this study was sourced from a range of latitudes exhibiting diverse climate and rainfall patterns representative of most of the species habitats (38.9-54.3°N, 116-128.7°W) and included 1382 genotypes total (**Figure 1A**). *P. trichocarpa* cuttings were received and planted in a greenhouse on February 1st, 2020. On March 30, 2020, they were relocated to a lath house to harden. Subsequently, 7,628 trees were transplanted to the field on April 10, 2020, with 782 unique genotypes in the drought field site and 1382 unique genotypes in the control field site, reflecting the availability of viable cuttings. A single row of *P. trichocarpa* border trees (not used for measurements) was established around the experimental trees in each replicate block to reduce the effect of undesired edge effects.

### Drought Treatment

We employed surface drip irrigation, applying a set water volume measured with flow meter sensors, OMNI™+ Turbo (T²) Water Meters, at each treatment site throughout the field season. Trees were fully irrigated until March 2021 to conclude their first establishment year. Following this, the irrigation was reduced in the drought treatment to achieve a soil moisture deficit of 150 centibars (0.15 MPa) from March to December 2021. This deficit was significant in comparison with the fully irrigated (control) treatment, representing a long-term modest drought on-going throughout the growing season. During the 2021 field season, soil moisture was monitored using Watermarks (Granular Matrix Sensors from MMM) (for daily soil water potential) and neutron probes (for volumetric water content every 12 days) across seven stations, with distinct placements for varying irrigation conditions. This comprehensive data, covering topsoil to a depth of 120 cm, verified an approximate 50% reduction in soil moisture for the drought treatment compared to the control, aiding our evaluation of drought’s impact on plant performance and water use efficiency **(Supplemental Figure S1)**. Soil moisture levels were systematically assessed during the 2021/2022 and 2022/2023 field seasons by using two primary techniques: 1) soil water potential was regularly recorded using Watermarks, equipped with remote daily data logging and 2) volumetric water content was determined using a neutron probe at 12-day intervals. A total of seven stations were established in the field: three within the drought site and four in the control site. At each station, two aluminum neutron probe access tubes (5 cm inner diameter) were deployed for soil moisture measurement. One tube was positioned in alignment with the irrigation line (indicative of a more moisture-rich site), while the other was placed between the rows of the irrigation line (approximately 100 cm away, representing a drier site distanced from the irrigation lines). For the watermark, seven sensors were installed at each of the seven drought monitoring stations mentioned above (a total of 49 sensors for all sites).

The neutron probe’s measurement depth reached a total of 120 centimeters. Additionally, at each of the seven drought monitoring stations, seven gypsum block watermark sensors were installed. The data collected from these sensors, encompassing both topsoil and deeper soil layers, were crucial in monitoring (as illustrated in Figure 3) and verifying that the areas under drought treatment exhibited an approximate 50% reduction in soil moisture compared to the control treatment. This significant difference in soil moisture levels was used to assess the impact of drought on crop performance and water use efficiency.

### Core population and sampling

We defined a subsample of unique genotypes that were fully replicated in triplicate in both drought and control treatments and this population is referred to as the "core population, n = ∼469 trees". However, note that some trees were lost due to mortality in the field **(Supplemental Table 1).**

For the leaf phenotyping, encompassing stomatal traits and leaf area measurements (**Table 1**), we conducted comprehensive sampling of the core population at two specific time points: June (one month after the drought treatment initiation) and September (106 days after the first collection, which includes an extended period of drought). In June, we sampled all of the core population of the drought treatment (Drought block 1,2,3 n= ∼3x469), and the core population control treatment we sampled only block 2 (n = 469). Similarly, in September we sampled the core population in blocks 1,2,3 (n = ∼3x469) of the drought and only block 2 (n= ∼469) in control, given time constraints for sampling and analysis. In summary, our analysis encompassed a total of 1,861 trees from both the June and September samplings, comprising 420 unique genotypes (441 trees) from the control group and 466 unique genotypes (1,420 trees) from the drought treatment.

Wood sampling for carbon isotope discrimination (**Table 1**) was performed on the whole “Core” population of all 6 blocks (469 genotypes, n = 3020 samples including replicates) in January 2022.

## Leaf Sampling

Two fully expanded, first and second mature leaves, specifically the fifth leaf from the apex on each of two south-facing primary branches, were collected at the height of approximately 1.3 m and, lightly misted with water, and sealed in plastic bags and stored in a cooler box on ice. These samples were immediately returned to the laboratory and subsequently kept at 4°C in a refrigerated room. All phenotypic analyses were initiated within 48 hours of collection.

### Stomatal Imprinting and Imaging

For the assessment of stomatal morphology, imprints were collected from the right abaxial portion of the leaf section and sampled as previously described (Tricker et al., 2004). A designated rectangular patch of this surface received a coating of clear nail polish. Upon drying, a strip of clear adhesive tape was utilized to lift off a thin layer, capturing a majority of epidermal cells and stomata. The tape was then placed on a microscopic slide and stored until further processing. This methodology resulted in a comprehensive collection of 1,869 imprints of all leaves collected **(example in Figure 1C**). The imaging phase involved using two Zeiss Standard 16x microscopes, each equipped with camera attachments. All imaging procedures used TCapture software (v5.1.1).

### Stomatal Detection and Analysis

To identify stomatal cells in our imprints, we integrated a model rooted in the U-Net architecture conceived within the PyTorch framework. The image data underwent a preliminary preprocessing stage and was transitioned into tensor format. Using a dataset of 300 representative images for training, our model was meticulously calibrated to distinguish stomatal cells in the imprints efficiently. During the predictive phase, the OpenCV, imported as cv2 (https://pypi.org/project/opencv-python/) software library facilitated the demarcation of the tightest bounding rectangle around every detected stomatal cell, an integral step for gauging its geometric characteristics. Our analytical process culminated to give the average stomatal dimension for each individual imprint **(example in Figure 1C**).

### Leaf area and perimeter

One of the two sampled leaves was used to measure leaf area (LA) and perimeter (PI). We used a Canon EOS Rebel T7i camera positioned over a lightbox (AGPTEK - Model A3USB) to photograph the leaves alongside a scale bar next to the leaves. Subsequently, these images were analyzed further at Oak Ridge National Laboratory (Oak Ridge, Tennessee). In this process, iterative thresholding was applied to the grayscaled image to separate the lightbox from the rest of the image, from which the leaf segmentation was extracted and rotated to achieve axis symmetry before being cropped to the leaf’s bounding box. Once the leaf was distinguished from the petiole, various leaf features, including leaf area and perimeter, were reliably detected and recorded **(example in Figure 1C**). The analysis was completed using the scipy (version 1.11.4) and scikit-image (version 0.19.3) libraries in Python.

### Dry weight and fresh weight

We measured the fresh weight of one of the two selected leaves, the same leaf that was photographed for the leaf perimeter, area, and fresh weight, using a precision balance (Adam Equipment - PGW 453e). Following this, the leaf was dried for 48 h in a paper bag, at 80°C and re-weighed for dry weight. To prevent moisture absorption, all dried leaves were stored with silica desiccant beads.

### Contact angle

Leaf contact angles were measured as described by (Kwon et al., 2014). A water droplet was placed on the left side of the leaf, whilst the right side was used for stomatal traits. To initiate the process, we prepared two leaf disks, which were subsequently taped to a microscope slide. For each leaf disk, a distilled water droplet was applied by a pipette. Subsequently, we captured images of the contact surface using a Canon EOS 500D camera. Similar to stomatal detection, we used a convolutional neural network implemented in PyTorch to distinguish water droplets following the region growing methodology (Lagergren et al., 2023). Using a dataset of 96 images for training and 23 for testing, we achieved a segmentation accuracy (Sørensen-Dice coefficient) of 0.917. Contact angle was measured by fitting a circle to the detected droplets using a Hough transform (Xu et al., 2013), and subsequently measuring the angle of the droplet to the microscope slide (Arnold 2021).

### Spectral Reflectance

Spectral reflectance was measured on the adaxial surface of each leaf utilizing a handheld PolyPen RP 410 (Photon Systems Instruments) after samples were brought back to the field. The data were subsequently captured and recorded using the SpectraPen software provided by the manufacturer. From the reflectance spectra, we computed NDVI, defined as Normalized Difference Vegetation Index **(see calculations in Table 1**), and PRI, defined as Photochemical Reflectance Index **(see calculations in Table 1**).

## Wood Sampling

### Preparation for δ^13^C

Wood samples for δ13C were collected in January of 2022 from the Core population in all 3 blocks of both treatments (control and drought). Sampling totaled 3019 samples and 479 genotypes, from which we harvested three 5 cm segments from the south-facing primary branches. These segments were carefully collected at a height of approximately 1.3 m and securely stored in labeled paper bags. Subsequently, the samples were dried at a constant temperature of 65°C, for a duration of 14 d.

Once dried, wood samples were extracted from the bags, and any remaining dormant buds at the branch tips were meticulously removed using shears. The samples were then ground to a fine powder, with a target of 0.50 grains of powder per milligram, utilizing a TissueLyser II (Qiagen) equipped with a Grinding Jar Set, Stainless Steel (2 x 10 ml) adapter. This grinding procedure involved a one-minute cycle at a frequency setting of 20 Hz, effectively reducing the dried wood to a powdered form similar to (Moghaddam et al., 2013) method. Subsequently, the ground wood samples were placed into labeled plastic vials, and precisely 3 mg (with a tolerance of +/-10%) was loaded into tin capsules and folded as needed for the analysis. Tin capsules were arranged in a 96-well plate, with two duplicates for each of the 33 well plates. These samples were then sent to the UC Davis Stable Isotope Facility for carbon isotope discrimination analysis using Isotope-Ratio Mass Spectrometry (IRMS).

### Calculations of δ13C

The following was used to calculate δ13C:

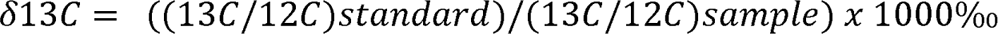

This value is expressed in parts per thousand (‰) and compares the ratio of 13C to 12C in the sample to that in a standard (typically a belemnite formation from the Peedee Formation in South Carolina, known as PDB). A negative δ^13C value implies that the ^13C/^12C ratio in the plant is lower than that in the standard. Since plants discriminate against d13C, most plants have negative δ^13C values. The more negative the value, the stronger the preference for ^12C in the photosynthetic process. A higher WUE is indicated by a higher ^13C/^12C ratio (less negative δ^13C values and less discrimination against ^13C).

## Tree height

Biomass productivity data were collected as tree height measurements in March 2021 (beginning of the growing season, after leaf flush), and November 2021 (end of growing season, after bud set), following the first year of the applied drought treatment. The height of every experimental tree across each block of both treatments was recorded in a team of 2 people. Using a telescopic height pole aligned with the base of the tree, one individual raised the pole to the same height as the shoot apical meristem of the dominant stem.

### Relative growth rate

The relative growth rate of height was calculated with this equation:

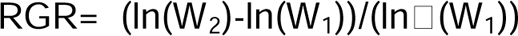

where:

W_1_ represents the height taken in March of 2021.

W_2_ represents the height measured in November of 2021.

## Statistical analysis

Statistical analyses were performed using RStudio (v4.1.2). We visualized spatial data using the Leaflet package (v1.7.1), a JavaScript library for interactive maps. After loading the necessary library, our geospatial data was overlaid on a CartoDB Positron basemap on which the data points can be placed. During the data exploration and organization phases, we employed a suite of packages including “dplyr”, “tidyverse”, “tidyr”, “plotrix”, “data.table”, and “stringr”. Enhanced data visualization was achieved with the “ggplot2” package, facilitating the creation of frequency diagrams, reaction norms, and boxplots. For statistical significance testing and Analysis of Variance (ANOVA), we utilized packages such as “stats”, “lme4”, “emmeans”, and “lmerTest”."

### Water content

The water content (WC) was also calculated by using (**see Table 1**):

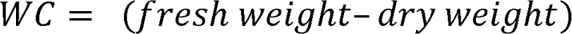

Proportional water content (PWC) was also calculated by using (**see Table 1**):

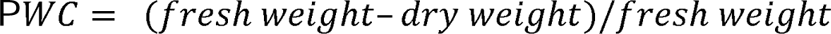

### Drought response

To quantify the plasticity of the trait, we employed the formula:

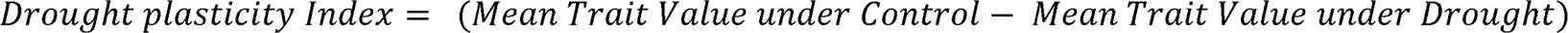

This index provided a relative difference value, representing the extent of trait variation due to drought exposure across different genotypes (**Figure 2B**).

**Figure 2.**
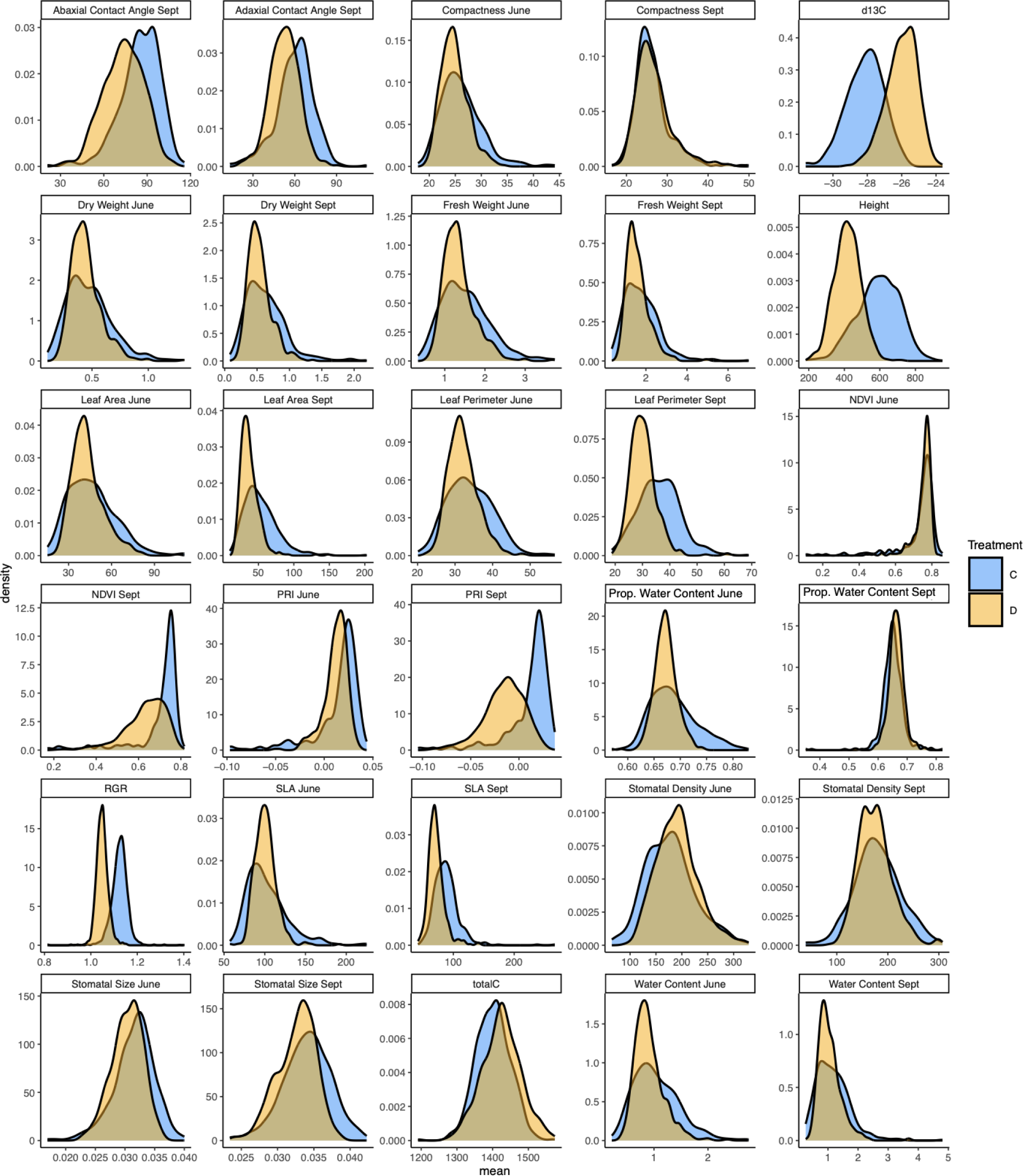
Density distribution comparisons for various traits under control (blue) and drought (orange) conditions.

### Drought Resilience index (DRI)

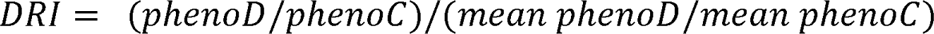

*phenoD* = Phenotype drought

*phenoC* = Phenotype control

*mean phenoD* = population mean of Phenotype D

*mean phenoC* = population mean of Phenotype C

This index calculates the ratio of yield reduction under stress in a specific genotype relative to the average reduction across all genotypes **(Supplemental Table 3).** PhenoD refers to a phenotype of the drought treatment and phenoC refers to a phenotype of the control treatment.

### Climatic data

Climatic data were sourced from WorldClim fit to western North America, which contains a set of global climate layers with high spatial resolution (2.5’), averaging across the years 1970-2000 (Fick & Hijman, 2017). The WorldClim database provides interpolated climate data (a total of 19 variables), including temperature, precipitation, and derived bioclimatic variables, facilitating a comprehensive understanding of the environmental climates of origins for our population of *P. trichocarpa* trees.

### Genotypic data

The reference genome and re-sequencing of the *P. trichocarpa* entire population (total of 1523 genotypes) has been conducted by the U.S. Department of Energy/Joint Genome Institute (Evans et al., 2014; Tuskan et al., 2006). The data can be accessed through the Joint Genome Institute Database. We used Genome version: *Populus trichocarpa v4.1, Phytozome genome ID: 533*.

### Pairwise trait correlations

To estimate the relationships between plant traits, pairwise Pearson correlation coefficients were calculated separately for control and drought treatments in R with cor(…, use=”complete.obs”). These correlations were visualized with a heatmap in R.

### Estimating variance components of traits

In our analysis, the contribution of Genotype (G), Treatment (T), and their interaction (G:T) to the variance in plant traits was estimated. This estimation was conducted in R, utilizing the lme4 package, where Genotype, Treatment, and Interaction were treated as random effects. Post-model fitting, the variance components were extracted using the VarCorr function, and the variance associated with Genotype, Treatment, Interaction, and Residual were isolated. The total variance was computed as the sum of these components. Subsequently, the percent variance explained by each component was calculated by dividing each variance component by the total variance and multiplying by 100.

Model:

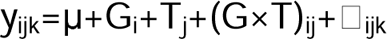

where:

y_ijk_ represents the observed trait value for the ith genotype, jth treatment, and kth observation.

μ is the overall mean of the trait across all genotypes and treatments.

G_i_ represents the random effect of the ith genotype.

T_j_ represents the random effect of the jth treatment.

(G×T)_ij_ is the random interaction effect between the ith genotype and jth treatment.

L_ijk_ denotes the residual error term associated with each observation, capturing the variability not explained by the other components in the model.

The percent variance explained by each component is calculated as:

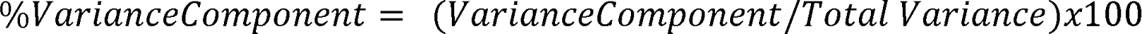

### Climate-Trait Correlations

For assessing relationships between traits (separately for drought and control treatments) and climate of origin (Bioclimatic variables 1-19 from WorldClim) of experimental trees, Spearman’s rank correlation coefficient was computed using R. The Spearman’s correlation measures the strength and direction of monotonic associations between paired data. This non-parametric test was chosen due to its robustness against outliers and its capability to detect non-linear relationships. Calculations were performed using the cor.test() function with the method set to "spearman’’ from the base R package.

### Random forest prediction of traits in relation to climate of origin

Utilizing the Random Forest package in R, we implemented a random forest regression model to analyze the impact of climate on plant traits (stomatal size, stomatal density, and δ13C) under drought conditions, informed by 19 WorldClim Bioclimatic variables. Our models, each comprising 500 trees, were validated to ensure convergence, confirming additional trees would not enhance predictive accuracy. After removing missing values, datasets were subjected to a complete case analysis. The model’s efficacy was visually assessed through scatter plots comparing observed and predicted trait values. Predictive mapping across diverse climates was conducted using bioclimatic data.

### Genome-wide association study (GWAS)

The Genome-wide association study (GWAS) was executed using the GEMMA software (Genome-wide Efficient Mixed-Model Association) to identify associations between the trait of interest and the genotypic data. GEMMA utilizes a linear mixed model approach to account for population structure and kinship, ensuring that the results are not confounded by underlying population stratification and has been used effectively for this population previously (Taylor et al., 2024). We filtered SNPs and Indels based on minor allele frequency = 0.05 (∼8 million after filtering). To identify potentially explanatory variants, we set a significance threshold of -log_10_(p) ≥ 5 for further consideration following the approach from (Blumstein & Hopkins, 2021). In cases where multiple variants passed this threshold and were within 400 kb of each other, we defined the “peak” as the most significant SNP within this region. To visualize candidate loci in manhattan plots, only peaks in which multiple variants passed our significance threshold were highlighted. Toward identifying potential candidate loci underlying these peaks we identified all genes within 20 kb of the peak region (spanning all variants passing the significance threshold for each peak). We used blastp (e<10^-5^) to identify similarity between proteins in these regions with *Arabidopsis thaliana* proteins (TAIR10) and checked the gene description and gene ontology annotations for “stomata”,”water”,”guard cell”, and “abscisic acid”. From this, we generated lists of loci, available in **Supplemental Table 6**, for future research aimed at identifying potential causal loci.

### Elucidating the Role of Chromosome 10 Locus in Stomatal Size

We examined, in greater detail, a locus on Chromosome 10 consistently associated with stomatal size under both drought and control treatments and at two measurement time points. The pronounced signal from this locus provided a unique opportunity for an in-depth case study, aiming to answer several pivotal questions:

To assess the LD across this genomic window, we calculated the Pearson correlation between all variant pairs in this 20 kb window, subsequently visualizing the results in a heatmap format and by visualizing LD with the most significant variant. Utilizing a random forest model (implemented in R using the randomForest package, with 500 trees), we modeled trait∼climate relationships based on genotypes with known phenotypic data. This model enabled us to predict stomatal size for genotypes with previously unmeasured traits based on their climate of origin, employing bioclimatic variables sourced from WorldClim 2.0. We then assessed the predicted allele effects by examining these predicted phenotypes with the respective allele states across our dataset Lastly, we investigated the relationship between allele state and climatic variables. This entailed calculating the t-statistic for the correlation between allele states and each individual bioclimatic variable. Furthermore, we employed a random forest model, incorporating all bioclimatic variables to predict allele states. The accuracy of this model was gauged through analysis of the confusion matrix in R. Predicted allele states were projected and mapped across the geographical range of *P. trichocarpa*.

### Future climate predictions of Chromosome 10 locus

To investigate the potential impact of future climate changes on the frequency of a specific allele associated with stomatal size on chromosome 10, we employed high-resolution future climate projections and predictive modeling. This allele was selected based on its significant association to stomatal size and with the climate of origin, suggesting its potential responsiveness to future climatic shifts. We utilized downscaled future climate projections (10-minute spatial resolution) from the Coupled Model Intercomparison Project Phase 6 (CMIP6), available through WorldClim version 2.1, which, as described (https://www.worldclim.org/data/cmip6/cmip6climate.html), underwent both downscaling and calibration (bias correction) against the baseline climate provided by WorldClim v2.1 Our analysis encompassed data from ten global climate models (GCMs), specifically: ACCESS-CM2, CMCC-ESM2, EC-Earth3-Veg, GISS-E2-1-G, INM-CM5-0, IPSL-CM6A-LR, MIROC6, MPI-ESM1-2-HR, MRI-ESM2-0, and UKESM1-0-LL. We evaluated these models across four Shared Socio-economic Pathways (SSPs): SSP1-2.6, SSP2-4.5, SSP3-7.0, and SSP5-8.5, at four future time intervals: 2021-2040, 2041-2060, 2061-2080, and 2081-2100.

Initially, we compared future climate predictions to current climatic conditions (baseline) using bioclimatic variables from WorldClimat the geographic locations of *P. trichocarpa* genotypes with t-tests, using T-test statistic as a unitless measure of the severity of change for each bioclimatic variable. This preliminary step established an understanding of the extent of future climatic changes potentially facing *P. trichocarpa*.

To predict future allele frequencies, we adopted a machine learning approach using random forest models in R. This involved analyzing the single nucleotide polymorphism (SNP) with the strongest correlation to stomatal size within the identified locus on Chromosome 10. Given the imbalanced genotype frequencies, we first balanced the dataset through random down-sampling. From this balanced dataset, we trained a random forest model to predict genotype frequencies based on bioclimatic variables. This model was applied to both the current climate baseline and all future climate scenarios, for each combination of GCM, SSP, and time interval.

To ensure robustness, this process was iterated 100 times with random down-sampling to obtain balanced training datasets, resulting in a total of 16,000 model predictions. For each prediction, we calculated the frequency of the allele associated with smaller stomata and compared it to the baseline frequency predicted from current climate conditions.

## Results

### Drought implementation

We measured variation in soil mean water potential at a 40 cm depth over the period from March 2021 to February 2022, which confirmed a water deficit in the drought treatment **(Supplemental Figure 1A)**. Upon implementing the drought treatment, we observed a significant decrease in the soil water potential, during the drought period. The water potential in the drought treatment (represented in orange) significantly decreased compared to the control treatment (represented in blue) over time, highlighting the successful implementation and effectiveness of the drought treatment in the field on soil water potential (**Supplemental Figure 1A)**. Additionally, through visual assessments, we identified signs of water deficit in *P. trichocarpa* trees, such as wilting or yellowing leaves and early leaf shedding. Trees in the drought treatment exhibited these signs more prominently than those in control treatment. **Supplemental Figure 1B)** presents the mean daily maximum and minimum temperatures, while **Supplemental Figure 1C)** displays precipitation data for Davis, California. Taken together, the stable soil temperatures and infrequent rainfall events suggest that the prevailing climate was particularly well-suited for our drought experiment.

### Genotype, Treatment, and Genotype X Treatment effects on leaf and stomatal traits

In brief, we measured 14 traits in replicates of ∼469 genotypes, in drought and control conditions at a field site in Davis, CA in July 2021 and September 2021 **(Materials and methods, Table 1**, **Supplemental Table 1, Supplemental Table 2, Supplemental Table 4)**.

The effect of drought relative to control conditions was evident in a number of traits measured, especially those related to biomass. Drought conditions significantly reduced leaf size, leaf mass, height, and growth rate (**Figures 2 and 3**). Among leaf morphological traits, we saw a trend for smaller more dense stomata under drought conditions, along with reductions in specific leaf area, leaf perimeter, and abaxial contact angle. This might reflect constraints on growth and whether the plasticity is adaptive remains a question for future work (**Figures 2 and 3**). Notably, water-related traits, notably δ^13^C, a proxy for water use efficiency, was significantly higher under drought, suggesting a water conservation response. For traits where measurements were made at multiple time points, the effect of drought was more pronounced on measurements made in September, after several months of water deficit, than in June, shortly after the onset of the drought treatment (**Figures 2 and 3**). This is especially clear in physiological measurements made via remote sensing. For example, the difference between control and drought for PRI and NDVI was most evident in September. In particular, PRI was dramatically reduced in drought-treated plants indicative of physiological stress (Magney et al., 2016; Mulero et al., 2023; Wong et al., 2022).

**Figure 3.**
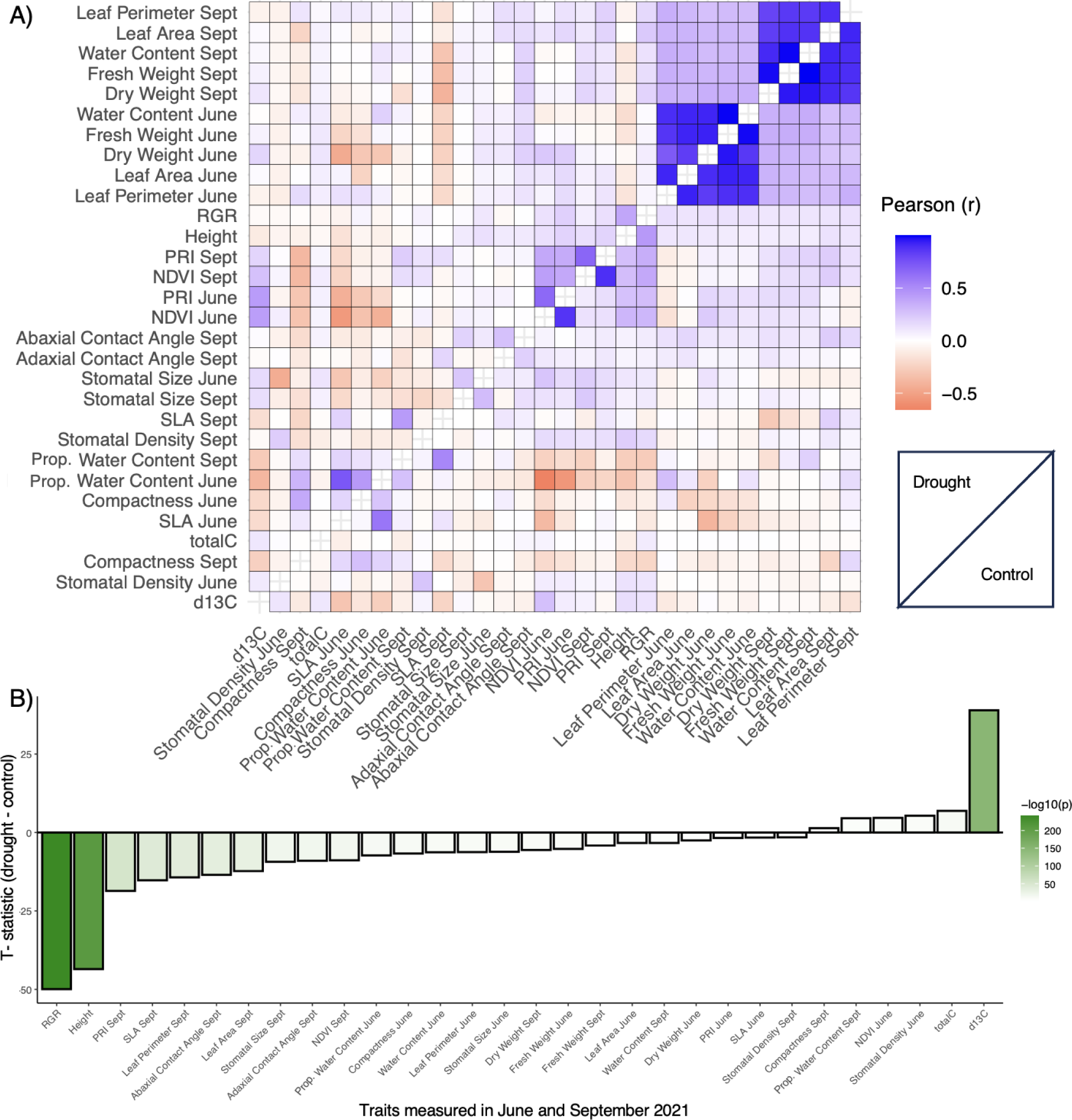
Comparative Analysis of Traits measured in this study. A) Correlations between traits. Above the diagonal are traits under drought conditions. Below the diagonal are traits measured under control conditions. B) Results of a pairwise T-test comparing genotype means between drought and control treatments for various plant traits of interest. Negative values indicate a trait reduction in drought conditions, and vice versa, with bars colored by the significance of effect (-log10(p)).

Trait correlations were similar under drought and control treatments (**Figure 3A**). Positive correlations were observed between Stomatal Size in June and Stomatal Size in September, indicating consistency in trait behavior across different growth stages and durations of drought (June vs. September). A negative correlation was evident between Stomatal Density and Stomatal Size, consistent with previous observations in poplar (McKown, Guy, Klápště, et al., 2014) and other species (Franks et al., 2009; C. Liu et al., 2021). We also observed significant plasticity in several traits. As expected, WUE (estimated with δ13C) increased under drought (**Figure 3B**). Other traits such as PRI, SLA, and stomatal size showed decreases under drought. However, it’s noteworthy that specific traits, namely Water Content, Stomatal Density, and Leaf compactness, demonstrate minimal deviation, suggesting their stability or marginal changes during drought conditions.

The variance components analysis from the experimental field trial delineated variability in plant traits into three sources: Genotype (G), Treatment (T), and their interaction (GxT). Genotypic variance was observed across most traits, underscoring the significance of genetic differences among the samples (**Figure S2A**). Environmental treatments significantly affected traits such as Stomatal Size in September and Leaf Compactness in June. The interaction between genotype and treatment was evident in traits such as Water Content, indicating a differential response of genotypes to various treatments. Weight-associated traits, including the Fresh Weight in September, displayed a balanced influence of both genetic and treatment factors. Overall, the findings emphasize the complex interplay of genetics and environment in determining plant trait variability, in particular, in response to drought. The PCA scatter plot **(Figure S2B)** reveals that both treatments, Control, and Drought, affect 14 leaf traits with some overlap, suggesting similarities in trait expression between treatments. The drought treatment exhibits a distinct shift along the first principal component, indicating a greater influence on the traits associated with PC1 (22% variance explained), while the Control treatment shows greater variability along PC2 (13% variance explained).

### Relationships between Traits and Climatic Conditions

We found discernible correlations between plant traits and the climate of their origin (**Figure 4**). Leaf size and weight measured in June and September, for instance, were predominantly associated with cooler, wetter climates, with larger heavier leaves associated with these cooler, wetter climates. Notably, stomatal density and size demonstrated contrasting relationships to climatic conditions. Specifically, genotypes originating from warmer, arid climates exhibited lower stomatal densities but possessed larger stomatal sizes. Under normal conditions, warmer, drier climates correlated with less negative δ13C values. However, this relationship was altered, and in some instances, even inverted when δ13C was measured under drought conditions, potentially suggesting the presence of adaptive plasticity in these genotypes (**Figure 4A**).

**Figure 4.**
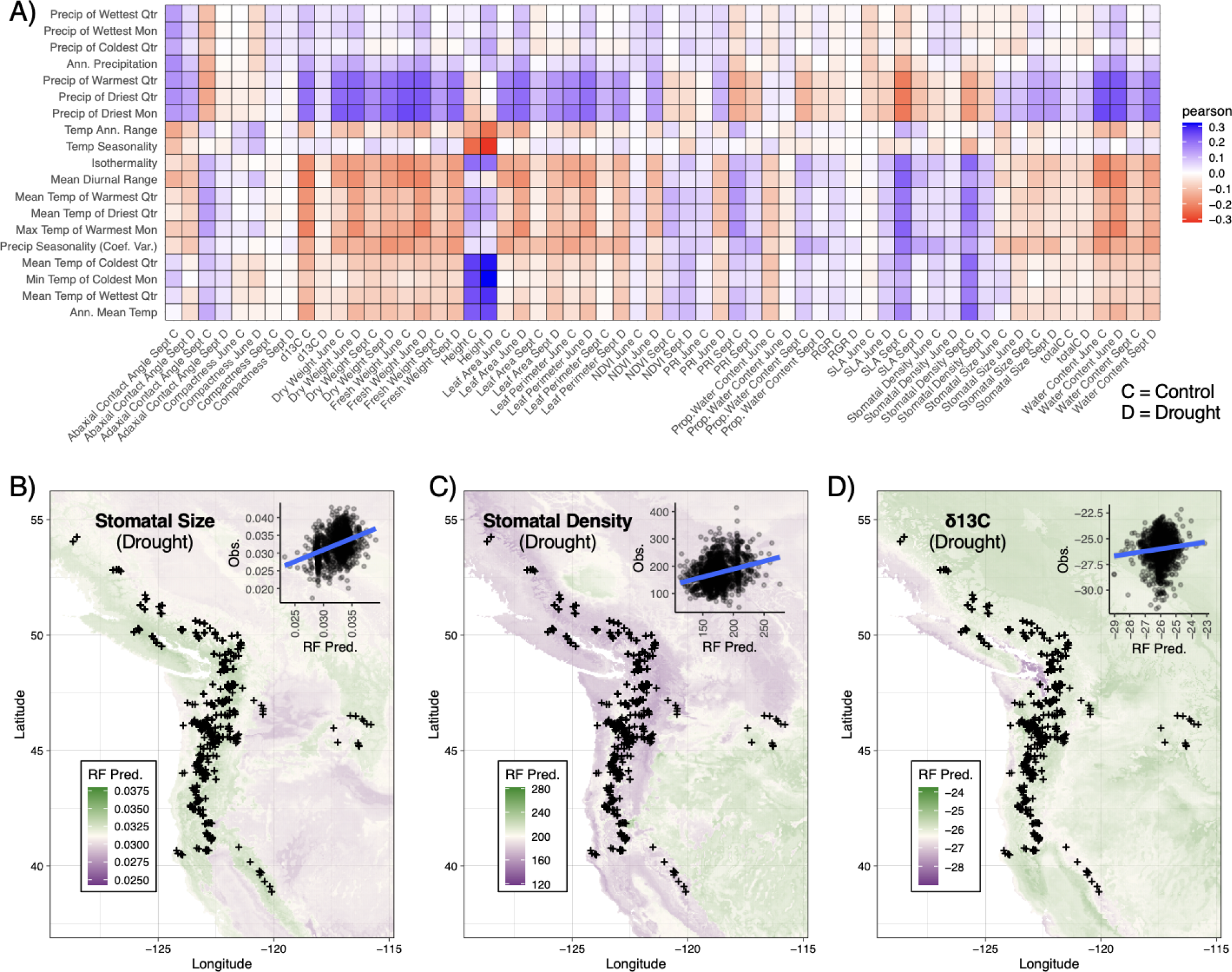
Analyses of traits in relation to climate-of-origin among *P. trichocarpa* genotypes. A) Heatmap illustrating Spearman correlations between stomatal traits and bioclimatic variables sourced from WorldClim. B-D) Geographic distribution of predicted B) stomatal size, C) stomatal density in September and D) δ13C under drought conditions based on Random Forest models, with scatter plot inset contrasting predicted versus observed values. In all geographic plots, the crosshairs indicate the locations of *P. trichocarpa* genotypes.

To provide a more nuanced understanding of the interaction between climatic variables and plant phenotypes, we employed Random Forest (RF) models (**Figure 4B-D**). The strength of RF models lies in their ability to detect non-linearities and interactions, thereby potentially enhancing the accuracy of predictions. Nevertheless, they do present challenges in terms of interpretability for targeted hypothesis testing. These RF models confirmed a marked correlation observed between the predicted values and the actual measurements. Visualizing these predictions across geographic landscapes further emphasized the negative correlation between stomatal size and density in response to climatic factors. For example, δ13C values under drought conditions were predicted to be lower in coastal regions, which are characterized by cooler temperatures, increased precipitation levels, and reduced climatic seasonality. Likewise, stomata were predicted to be smaller and more dense in cooler, wetter environments.

### Genome-Wide Associations (GWAS)

We performed three GWAS analyses for each trait: phenotypes under drought, control, and the difference (plasticity) (**Figure 5**). These analyses identified numerous loci explaining variance in traits after accounting for population structure. In total, across all traits, environments, and plasticity, we identified 1610 peaks that contained at least 3 variants that reached a significance threshold of - log_10_(p)≥5, a liberal threshold chosen to prioritize the trade-off between type I and type II errors and prevent false negative associations. To further refine results and identify potential candidate loci, we further localized genes to GWAS peaks and used blastp to identify potential orthologs in *Arabidopsis thaliana* with annotated functions. These analyses revealed many genes with annotations (e.g., Gene Ontology) with functions relating to water response, abscisic acid, guard cells, and stomatal development, though further work is necessary to test candidate loci. Given the large scope of these results, they are presented in total in **Supplementary Tables 4-5**.

**Figure 5.**
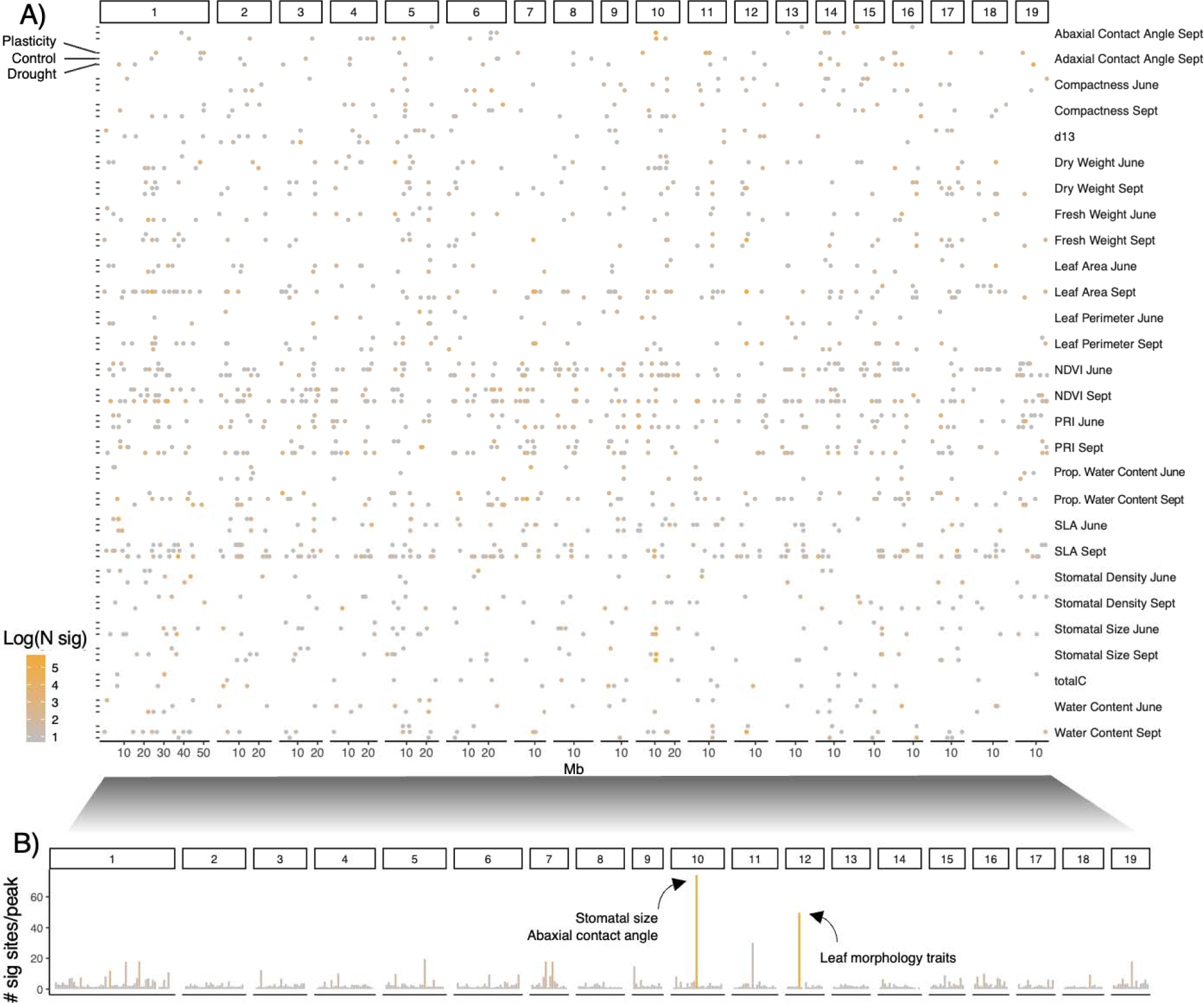
Summary of Genome-Wide Association Study (GWAS) results across traits. A) Points show the position of peaks identified, colored by the log(number) of variants with -log10(p) ≤ 5 across multiple traits along the *P. trichocarpa* Genome. Each dot corresponds to a significant genetic association for the designated trait at the specific genomic location. B) Meta-plot showing total number of variants with -log10(p) ≤ 5 across all traits, to identify hotspots potentially associated with multiple traits.

Summarizing GWAS results across traits revealed a notable locus on Chromosome 10. This region was significantly associated with stomatal size in both drought and control treatments, and for measurements in both June and September, as well as for abaxial contact angle, a proxy measure of cuticle wax (**Figure 5**, **Supplemental Figure S3)**. We found that alleles in this locus explained 15.9% of the variance in stomatal size across all measured plants, irrespective of treatment. Spanning over 200 kb and encompassing numerous genes, it is challenging to pinpoint a singular causal variant (**Supplemental Figure S3**). It is also plausible that the peak might represent a series of closely associated causal variants. Interestingly, this region contained a cassette of 11 tandem 3-ketoacyl-CoA synthase (KCS) family genes, with KCS genes being previously linked to both cuticular wax and stomatal traits (**Supplementary Tables 5)** (Gonzales-Vigil et al., 2017; Gray et al., 2000). Indeed, a KCS gene in this region was previously linked to high alkene content in abaxial cuticular wax and differences in stomatal size in *P. trichocarpa (Gonzales-Vigil et al., 2017)*. The linkage disequilibrium around this region revealed that the variants with strong associations had significant LD with one another, consistent with a solitary causal locus with extensive LD with neighboring variants **(Supplemental Figure S3**). Further analysis of allele effects across both environments demonstrated a consistent directionality throughout this window, where variants correlated with smaller stomatal size, though the existence of several linked alleles with the same effect direction in this region cannot be ruled out.

To explore its role in local adaptation to climate, we estimated allele effects on climate-predicted phenotypes for *P. trichocarpa* genotypes whose stomatal phenotypes were not analyzed in this study (**Figure 6**). The rationale behind this approach was multi-fold. Primarily, it served as a strategy to verify the integrity of climate-trait associations. If the phenotype predictions based on climate were accurate and reflect local adaptation, they should replicate the empirical allele effects observed in genotypes for which phenotypes were measured. Secondly, it showcases the potential of leveraging local adaptation in employing climate-predicted phenotypes in a broader genotype sample to augment efforts in pinpointing physiologically significant loci. These analyses are comparable to environmental GWAS tests which treat environment of origin as a phenotype, e.g., (Ferrero-Serrano & Assmann, 2019), but instead capture the environmental parameters predictive of a target trait, thereby potentially enhancing the study of locally adaptive alleles.

**Figure 6.**
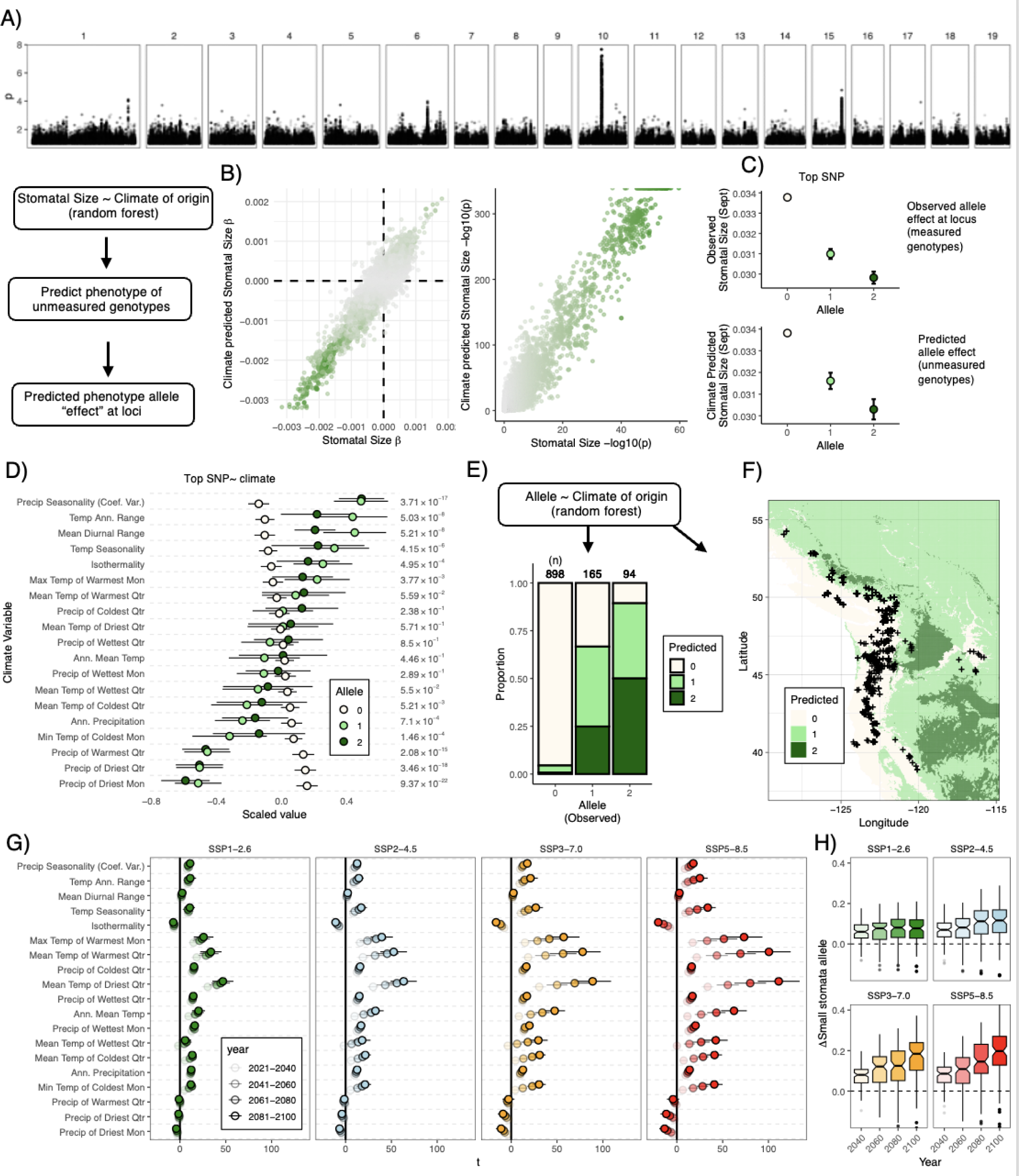
Analysis of Locus on Chromosome 10 (10,356,950 – 10,746,695) associated with Stomatal Size. A) GWAS of Stomatal Size summarized (mean -log10(p)) across both treatments and both times measured (June and September). B) left: Allele effects on stomatal size (left:β, right: -log10(p) from simple linear regression phenotype∼allele across all observations and ”allele effects” on climate-predicted allele effects of genotypes with unmeasured phenotypes in region on chromosome 10. C) Depiction of allele effect on stomatal size for genotypes with measured (upper) and climate-predicted (lower) phenotypes. Each point illustrates the mean, while error bars (barely visible) represent ± 2 standard errors. D) T-Statistic values from Pearson Correlation, illustrating the relationship between allele state and the climate origin of genotypes. E) Visualization of the prediction accuracy of the random forest model, presenting the proportion of predicted allele states corresponding to each observed allele. F) Geographical mapping of predicted allele states based on bioclimatic variables. G) change in bioclimatic variables from historical measures under various climate scenarios (SSPs and time). X-axis (T) represents t-test difference between future and current measures. Points mark mean values, where error bars indicate standard error across 10 GCM models. I) predicted change in frequency of small stomata allele on Chromosome 10 under future climates. Boxplots represent predictions averaged across GCM models for 100 iterations of randomized balanced training sets for random forest models of genotype ∼ bioclim variables under contemporary climates, used to predict genotype states at *P. trichocarpa*.

These analyses confirmed that observed allele effects were highly predictive of allele effects on phenotypes predicted by climate of origin in unmeasured genotypes (stomatal size: r = 0.95, p<2x10^-16^). Applying this methodology to the most significant SNP within this region yielded two noteworthy findings: an observable additive effect where heterozygous individuals presented an intermediate phenotype and a match between the empirical "effect" and climate-predicted phenotypes (**Figure 6**). Further exploration into climate-allele associations revealed that alleles linked with small stomata were predominant in environments characterized by reduced warm-season precipitation and greater precipitation seasonality (**Figure 6D**, **Figure S4)**. This aligns with the general trait correlations we noted, consistent with the distribution of alleles at this locus reflecting local adaptation to climate. The robustness of the relationship between the climate and allele state is further emphasized when predicting the allele state using the random forest model based on climate parameters (X^2^ test, p<2x10^-16^, **Figure 6**).

### Future climates

Given the strong link between loci affecting stomata size and climate, we set out to ask whether this yields predictions about future evolution at this locus. We evaluated the severity of climate change across various bioclimatic variables at the geographic locations of *P. trichocarpa.* These were based on different Shared Socio-economic Pathways (SSPs) and temporal projections, revealing that more intense SSP scenarios forecasted more substantial changes in climate, intensifying over time (**Figure 6G and I**).

As expected, the predicted changes in climate suggest a general trend toward warming. Notably, there is an expected increase in the extremities of precipitation - wetter conditions during the colder months coupled with drier conditions throughout the summer months. These shifts in precipitation patterns may drive natural selection on stomatal size. The allele associated with smaller stomatal size is found to be more prevalent in environments with high intra-annual precipitation variability and drier summer months—conditions that are expected to become more common in the future for *P. trichocarpa* (**Figure 6G and 6H**).

Our predictive models, averaging across all 10 Global Climate Models (GCMs) for each set of 100 random iterations, indicated a significant increase in the expected frequency of the small stomata allele, particularly under the more severe climate scenarios (SSP3-7.0 and SSP5-8.5), with a notable escalation over time (**Figure 6G and 6H**). These increments were most pronounced in the latter half of the 21st century. Taken together, these findings support the hypothesis that allelic variation associated with stomatal size reflects local adaptation to climate, which will be perturbed by extreme climate change.

## Discussion

This study has identified novel genetic loci that underpin phenotypic variation in stomatal and other leaf traits that are potentially important in determining *P. trichocarpa* crop adaptation to future water-limited environments. These findings will be informative in enabling future feedstock development for marginal environments. By deploying a controlled drought experiment, subjecting over 7,000 trees across 6.1 ha to a controlled soil moisture deficit we have begun to unravel the complexities of adaptive versus plastic responses to drought.

Our analysis indicates that stomatal traits are closely linked to the climatic conditions of their origin, with genotypes from hotter, drier climates often featuring smaller, denser stomata and enhanced water use efficiency. This relationship is valuable for considering poplar as a bioenergy crop in marginal lands and could help inform the selection of genotypes with stomatal traits and other traits aligned to the target environment of production. Consistent with previous findings (Pearce et al., 2006), our results affirm that trees adapted to dry, warm climates develop small, dense stomata to optimize water use efficiency under water scarcity, an essential evolutionary adaptation for maintaining gas exchange and water conservation in stressful environments. Furthermore, the plastic response of stomatal size reduction under drought conditions suggests an adaptive advantage, supporting smaller stomata associated with arid environments and underscoring their role in optimizing water use efficiency (Dunlap & Stettler, 2001). No strong relationship between stomatal size and height or RGR (**Figure 3**) was observed, supporting the idea that these traits can be disentangled enabling future selection and breeding of adaptive stomatal traits for hot and dry environments, with maintained yield. While WUE increased under drought, potentially reflecting adaptive plasticity, the relationship with stomata size was not evident.

A significant locus on Chromosome 10 was identified, associated with stomatal size, showing a distribution among genotypes highly predicted by the climate of origin. This is interesting since a significant locus on chromosome 10 was previously found to be in determining δ13C in a bi-parently mapping population with one grandparent as *P. trichocarpa* (Viger et al., 2013), thus linked to plant WUE. This finding is consistent with a scenario of local adaptation and is supported by additional loci identified through GWAS for multiple traits. Of the known candidates that map to this Chromosome 10 locus, 3-Ketoacyl-COA synthase 11, KCS11 is one such candidate that is a gene family member involved in the biosynthesis of Very Long Chain Fatty Acids (VLCFAs) codes for long-chain fatty acids, already shown to have a role in stomatal development and function in relation to drought tolerance and stomatal response to elevated carbon dioxide (Gray et al., 2000; Tang et al., 2020). These genetic markers provide a valuable resource for future studies aiming to dissect the genetic basis of adaptation in *P. trichocarpa*. In addition to adaptation, we identified significant plasticity in multiple traits, associations between phenotypes and climate of origin, and genetic loci underlying genetic variation. Our findings significantly extend previous work in poplar suggesting that phenotypic traits can be significantly influenced by geographical and environmental gradients (McKown et al., 2019; McKown, Guy, Klápště, et al., 2014). Here we subjected a large wild collection of *P. trichocarpa* to a controlled drought enabling us to investigate plastic versus adaptive variation. Northern and coastal genotypes are generally characterized by larger stomatal pores, faster growth rates, better stomatal conductance, higher carbon gain, and lower WUE (McKown et al., 2019; McKown, Guy, Klápště, et al., 2014; McKown, Guy, Quamme, et al., 2014). Our findings provide compelling evidence for local adaptation in *P. trichocarpa*, with clear associations between climate variables and key functional traits (Viger et al., 2016).

*P. trichocarpa*’s current adaptation to its climate underlines the need to explore resilience under changing future climates, since the ability to predict alleles with future adaptive value is essential for conservation and the management of natural populations (Blumstein et al., 2020). The adaptability shown by *Populus balsamifera*, through a wide range of adaptive physiological responses, offers hope for potential resilience against future climate shifts (Keller et al., 2011). The clinal variation and genomic signals observed in *P. trichocarpa* indicate a blend of unique and shared adaptive responses to different environmental gradients, shaped by genomic, environmental, and functional factors (Zhang et al., 2019). Such insights lay a groundwork for understanding adaptation mechanisms in species facing climate change threats.

We found novel evidence of predicted allele frequency shifts towards those associated with smaller stomatal size under future climate projections, especially under severe SSP3-7.0 and SSP5-8.5 scenarios, suggesting that natural selection may favor traits conducive to surviving anticipated shifts in precipitation patterns (**Figure 6**). This trend prompts crucial considerations about the pace of evolutionary dynamics in perennial species like *P. trichocarpa* in response to forecasted rapid environmental changes and the implications for the species’ persistence and resilience. In contrast, for research on conifer trees, there is evidence of limited adaptation to local climate for populations of Douglas Fir (Candido-Ribeiro & Aitken, 2024) although this study largely considered photosynthetic traits only and is in contrast to earlier research on Douglas Fir that demonstrated significant intraspecific population variation in traits in relation to climate of origin (Bansal et al., 2015), or relevance to identifying adaptive and plastic responses to enable drought tolerance in the face of climate change. For *P. trichocarpa,* our data suggest significant potential for allele frequency shifts with climate, but the timing and importance of these phenomena require further investigation.

The locus identified on Chromosome 10, related to stomatal size, emerges as a promising candidate for genetic exploration to enable precise gene editing (Allwright & Taylor, 2016; Taylor et al., 2019, 2024). Investigating this and other loci could reveal causal variants, providing breeders with tools to develop *Populus* varieties tailored for specific environmental demands, thus enhancing productivity and stress resilience. This approach aligns with sustainable biofuel production goals and broader environmental stewardship efforts. However, detailed mapping of these regions is necessary to unravel underlying genetic mechanisms (**Supplemental Table 6**).

Furthermore, predicting stomatal size from the climate of origin offers a novel approach to inferring phenotype values in genotypes without direct measurements, aiding in the association between observed alleles and traits. This methodology holds potential for streamlining the assessment of complex physiological traits, though its applicability across various traits and species requires more empirical validation.

The potential of poplar as a fast-growing short-rotation future feedstock for Sustainable Aviation Fuel (SAF) production is well-known (Porth & El-Kassaby, 2015; Sannigrahi et al., 2010) and underscores the importance of our research to underpin feedstock breeding programs to enhance crop efficiency, yield, and adaptability under current and future climate scenarios, particularly for marginal, water-limited sites, acknowledged as a limitation for widescale bioenergy crop deployment (King et al., 2013). Utilizing natural poplar populations for breeding can help preserve unique traits and biodiversity, preventing trait loss through domestication or selective breeding, and can help to reduce the long breeding cycles traditionally associated with forest tree improvement, since the complex allele-phenotype relationships can be preserved and investigated. Marker-assisted selection, based on single markers, has often struggled to deliver for tree improvement, given the high complexity of traits. In contrast, population-based approaches using Genomic Selection, where genome-wide markers are used to predict traits, particularly outside of the known climatic envelope, such as described here, offer significant future potential, to inform and refine breeding strategies for improved woody feedstocks, as has been demonstrated recently for other forest trees (Grattapaglia et al., 2018)( and is highly applicable to adaptive breeding in the face of a changing climate (Cortés et al., 2020; Depardieu et al., 2020). In addition to this, the research described here, also has significant potential, as appropriate, to deliver single gene targets for gene editing, already demonstrated in *Populus (Zhou et al., 2015)* and of value for accelerated wood domestication in forest trees (Anders et al., 2023). Taken together, our research provides multiple opportunities to develop improved woody feedstocks, optimized for future climates, that will contribute significantly in enabling the US to effectively triple the supply of biomass, contributing to a 15 % of energy demand, with dedicated energy crops, an important part of the mix of biomass supply, aligning with broader goals of energy sustainability and environmental conservation (U.S. Department of Energy, 2024).

By addressing the outlined gaps and expanding our research scope, we can contribute to a more nuanced understanding of the dynamic interaction between climatic factors and plant trait variation. This knowledge is crucial for informing more effective conservation strategies and breeding programs, paving the way for the sustainable use of poplar and other species in a changing global climate.

## Conclusion

We have uncovered evidence of adaptive variation in stomatal and leaf traits within *P. trichocarpa*, pinpointing a set of loci associated with genetic variation under both drought and control conditions, as well as their plasticity. Our findings also underscore the critical role of climate in shaping the adaptive landscape of *P. trichocarpa*, casting light on potential challenges and opportunities for this species in the face of climate change. Leveraging climate-predicted phenotypes provided compelling evidence for the adaptive significance of a locus associated with stomata size, which is predicted to respond to selection under future climates. In summary, this work offers pivotal insights into the intricate genetic underpinnings of physiological traits and their drought responses in *P. trichocarpa,* with implications for both conservation strategies and breeding initiatives.

## Abbreviations

CID: Carbon Isotope Discrimination
CO_2_: Carbon dioxide
SLA: Specific leaf area (Leaf surface area [mm^2^]/ dry mass [mg])
δ^13^C: ratio of the two stable isotopes of carbon ^13^C and ^12^C, a proxy for water use efficiency
GWAS: Genome-wide association study
NDVI: Normalized difference vegetation index
PRI: Photochemical Reflectance Index
RGR: Relative Growth Rate
PWC: Proportional Water Content
SAF: sustainable aviation fuel
WUE: Water-use efficiency

## Acknowledgments

This research was conducted on the ancestral land of the Patwin people. University of California, Davis land acknowledgment statement: https://diversity.ucdavis.edu/land-acknowledgement-statement Furthermore, we thank all UC Davis interns, graduate students, and postdocs, who helped with field collections and lab assistance on this project.

## Data availability

Phenotypic data and GWAS results are provided as supplemental files. Sequencing data are available on the NCBI SRA (PRJNA247507 -PRJNA256895). The Populus trichocarpa v4.1 JGI reference genome is available on Phytozome 13. Worldclim climatic variables are available from https://www.worldclim.org/.

## Code availability

https://github.com/mchklein/Poplar-Stomata

## Supplemental Materials

### Supplemental Materials and Methods

NA.

### Supplemental figures

**Figure S1. Soil water potential and Davis, CA climate data.**

A) Soil mean water potential at 40 cm depth. Data derived from gypsum block water marks installed across drought and control treatments (control = blue; drought = orange) field site from March 2021 until February 2022. The ribbon around the graph line represents standard error. In B) the Mean, Max and Min daily temperature and C) Daily precipitation from Davis, California from March 2021 until January 2022 are shown. Data from: “DAVIS 2 WSW EXPERIMENTAL FARM, CA US USC00042294’’ www.ncei.noaa.gov.

**Figure S2. Variance components analysis A) and Principal Component Analysis B) by trait.**

A) The stacked bar chart represents the distribution of variance components for stomatal and leaf plant traits, assessed in two months: June and September. Variance sources are color-coded: genetic (G) in blue, environmental (T) in green, and genotype by environment interaction (GxE) in red. The x-axis displays the percentage variance explained, while the y-axis lists the specific traits under investigation. For several traits, the GxE interaction explains a substantial proportion of the observed variance, emphasizing the importance of considering both genetic and environmental factors in trait expression.

B) Principal Component Analysis (PCA) Scatter Plot of 14 Different Leaf Traits visualizing the variation of 14 different leaf traits between the two treatments, control and drought. The horizontal axis, PC1, explains 26% of the variance, while the vertical axis, PC2, accounts for 12% of the variance.

**Figure S3. Genome-Wide Association Study (GWAS) results for stomatal size.** A) Stomatal Size in June under control (upper) and drought conditions (lower) and Stomatal Size in September control (upper) and drought conditions (lower). A prominent peak is observed on Chromosome 10 across both environmental conditions and time points. This peak is delineated by the green boxes and further detailed in the zoomed-in panels on the right, which center on the most significant variant. Green icons within the zoomed-in panels mark the positions of *P. trichocarpa* genes. The most significant polymorphisms for each trait are emphasized in the respective panels. B-D): Effects and linkage disequilibrium locus on Chromosome 10 (10,356,950 – 10,746,695) associated with Stomatal Size. B) Estimated genetic effect (β and p from simple linear regression phenotype∼allele across all observations) on size and density for each variant in this window. C). Estimated LD (Pearson correlation) between all variants and the variant with most significant association to each trait. D). Relationship between LD of each variant with the most significant variant and the significance of their association to each trait colored by their estimated genetic effect (β).

**Figure S4. Observed geographical distribution of example allele in Chromosome 10 (10,356,950 – 10,746,695) associated with stomata size and density.**

### Supplemental Tables

**Supplemental Table 1. Raw phenotypes.**

**Supplemental Table 2.**
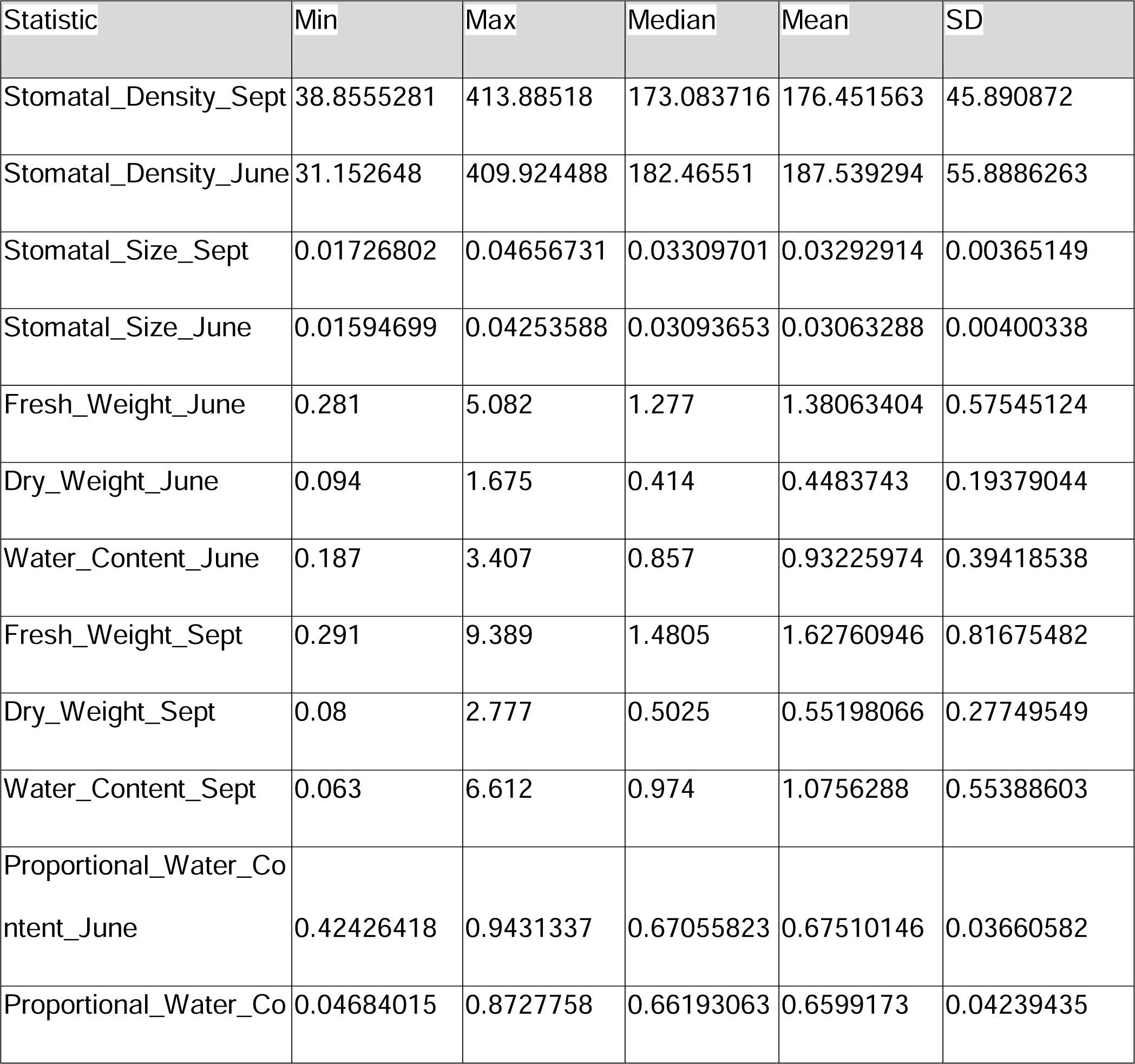
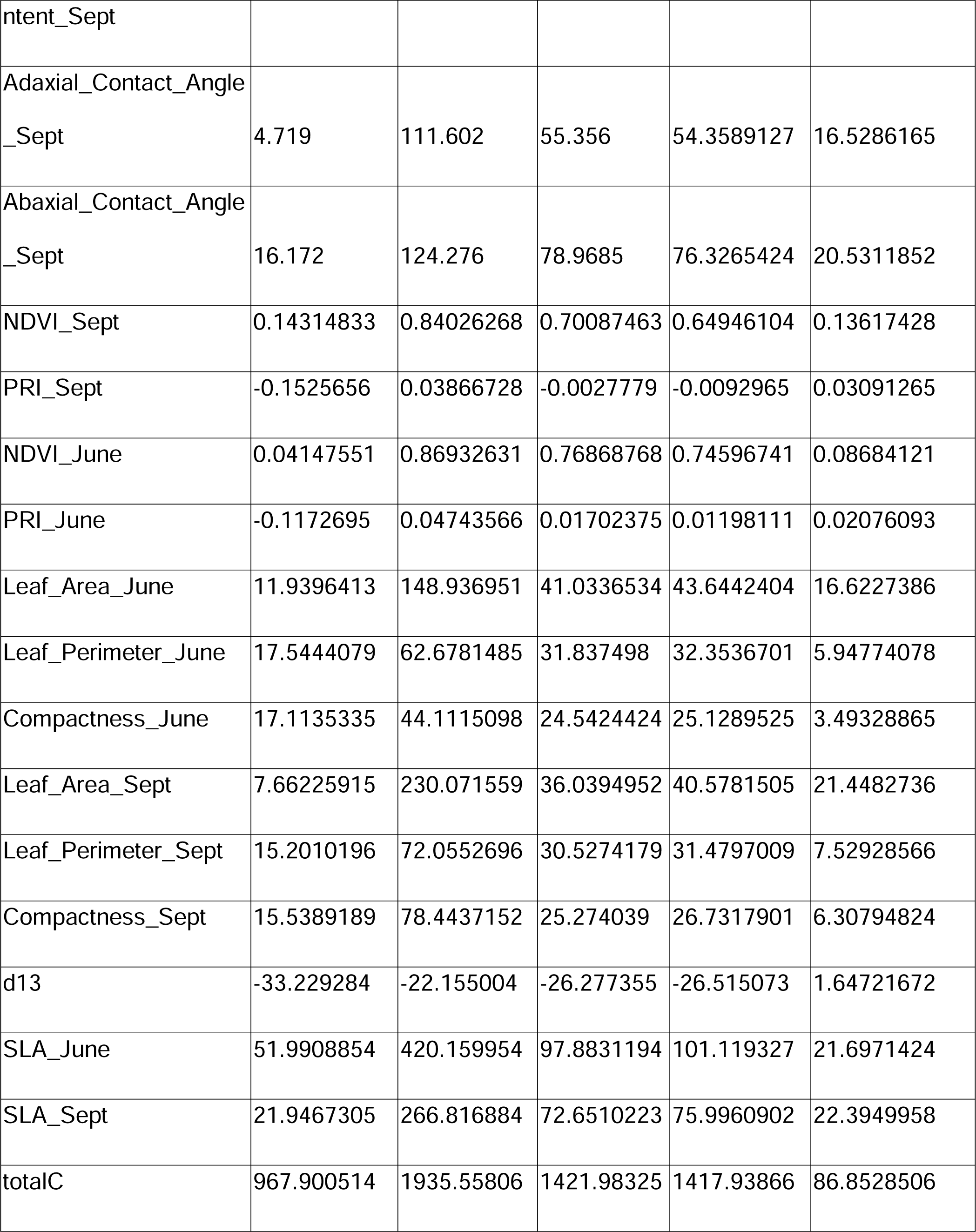
(Min/max/average) for each phenotype measured.

**Supplemental Table 3. Drought Recovery Index (DRI)**

**Supplemental Table 4. Genotype information (location, climate).**

**Supplemental Table 5. GWAS summary**

**Supplemental Table 6. All Potential Candidate genes (Orthologs to *Arabidopsis thaliana*), also highlighted with 4 categories (Water, Stomata, Guard Cells, ABA)**

