## Supplemental Figure 4 for "Climate adaptation in *P. trichocarpa*: key adaptive loci identified for stomata and leaf traits"

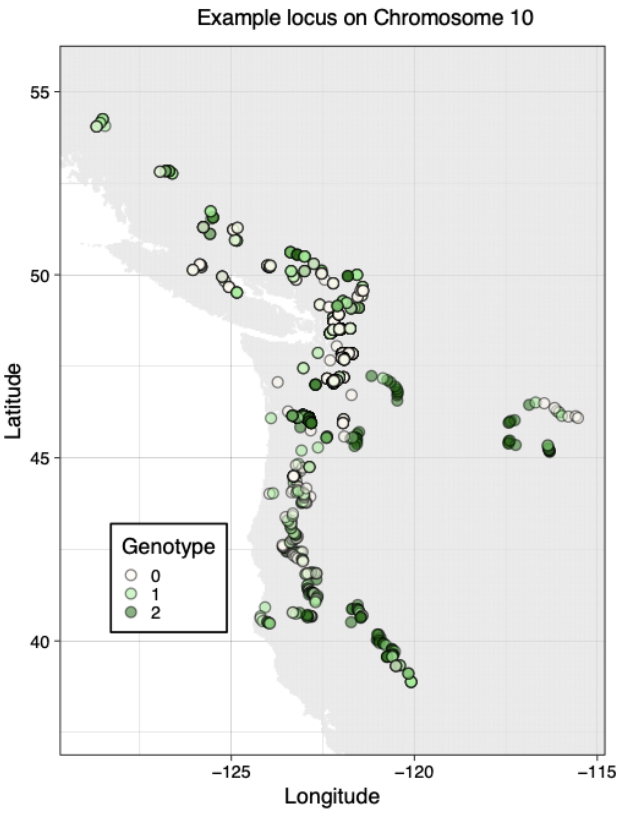


**Figure S4. Observed geographical distribution of example allele in Chromosome 10 (10,356,950 – 10,746,695) associated with stomata size and density.**
