## Supplemental Figure 3 for "Climate adaptation in *P. trichocarpa*: key adaptive loci identified for stomata and leaf traits"

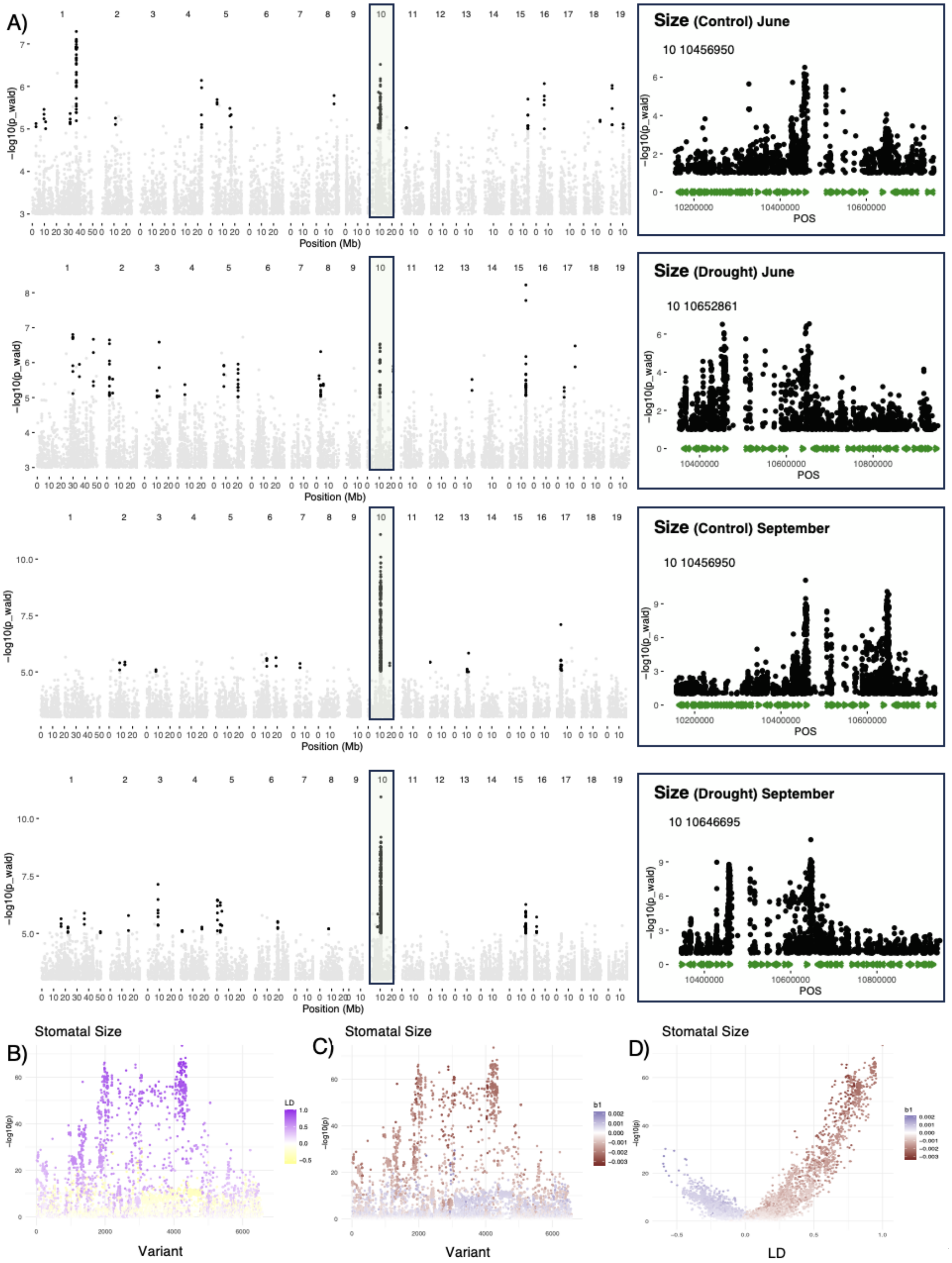
A)
Position (Mb)
−log10(p_wald)
Size (Control) June
10 10456950
Size (Drought) June
10 10652861
Size (Control) September
10 10456950
Size (Drought) September
10 10646695
POS
B)
Stomatal Size
Variant
LD
C)
Stomatal Size
Variant
b1
D)
Stomatal Size
LD
−log10(p)

**Figure S3. Genome-Wide Association Study (GWAS) results for stomatal size.** A) Stomatal Size in June under control (upper) and drought conditions (lower) and Stomatal Size in September control (upper) and drought conditions (lower). A prominent peak is observed on Chromosome 10 across both environmental conditions and time points. This peak is delineated by the green boxes and further detailed in the zoomed-in panels on the right, which center on the most significant variant. Green icons within the zoomed-in panels mark the positions of *P. trichocarpa* genes. The most significant polymorphisms for each trait are emphasized in the respective panels. B-D): Effects and linkage disequilibrium locus on Chromosome 10 (10,356,950 – 10,746,695) associated with Stomatal Size. B) Estimated genetic effect (β and p from simple linear regression phenotype~allele across all observations) on size and density for each variant in this window. C). Estimated LD (Pearson correlation) between all variants and the variant with most significant association to each trait. D). Relationship between LD of each variant with the most significant variant and the significance of their association to each trait colored by their estimated genetic effect (β).
