## Supplemental Figure 2 for "Climate adaptation in *P. trichocarpa*: key adaptive loci identified for stomata and leaf traits"

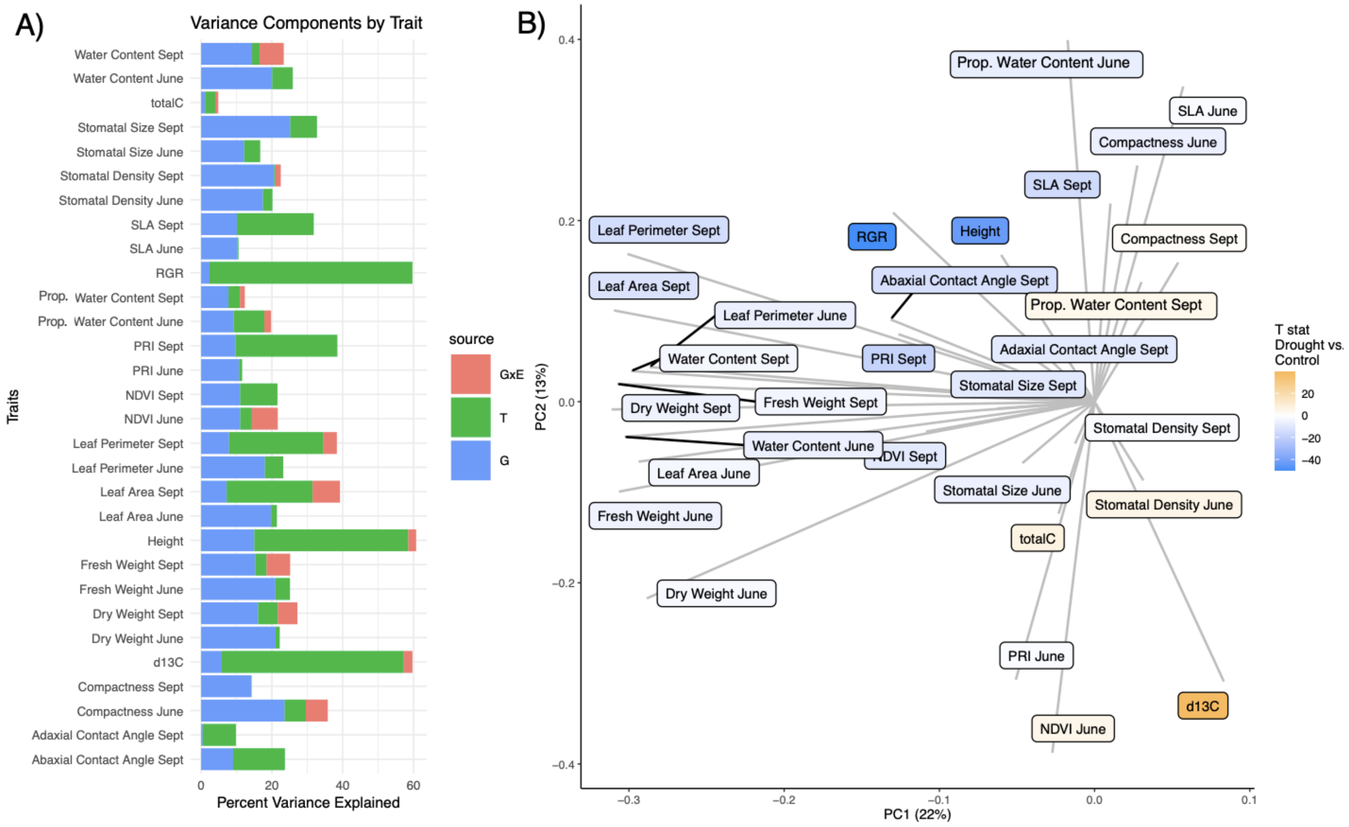


**Figure S2. Variance components analysis A) and Principal Component Analysis B) by trait.** A)The stacked bar chart represents the distribution of variance components for stomatal and leaf plant traits, assessed in two months: June and September. Variance sources are color-coded: genetic (G) in blue, environmental (T) in green, and genotype by environment interaction (GxE) in red. The x-axis displays the percentage variance explained, while the y-axis lists the specific traits under investigation. For several traits, the GxE interaction explains a substantial proportion of the observed variance, emphasizing the importance of considering both genetic and environmental factors in trait expression. B) Principal Component Analysis (PCA) Scatter Plot of 14 Different Leaf Traits visualizing the variation of 14 different leaf traits between the two treatments, control and drought. The horizontal axis, PC1, explains 26% of the variance, while the vertical axis, PC2, accounts for 12% of the variance.
