## Supplemental Figure 1 for "Climate adaptation in *P. trichocarpa*: key adaptive loci identified for stomata and leaf traits"

### Supplemental figures

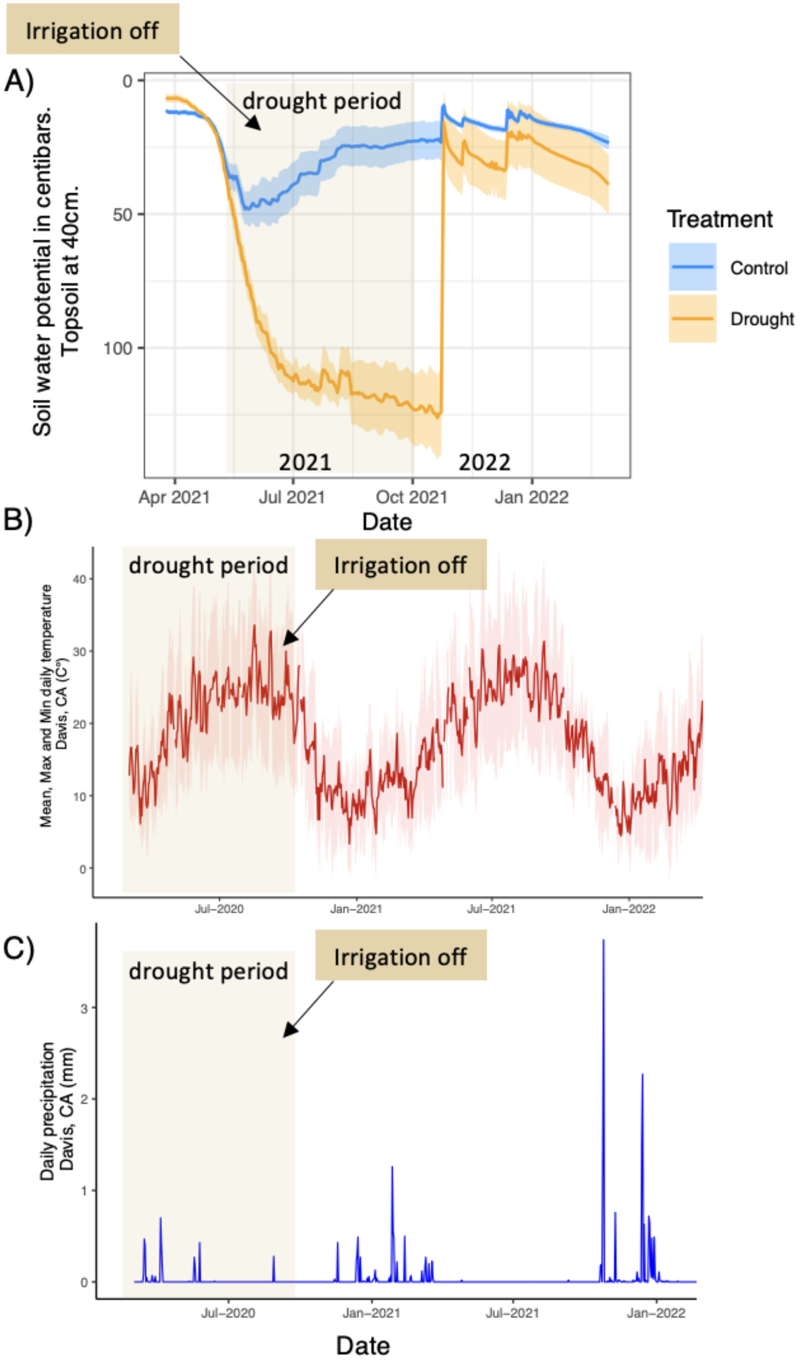

**Figure S1. Soil water potential and Davis, CA climate data.**

A) Soil mean water potential at 40 cm depth. Data derived from gypsum block water marks installed across drought and control treatments (control = blue; drought = orange) field site from March 2021 until February 2022. The ribbon around the graph line represents standard error. In B) the Mean, Max and Min daily temperature and C) Daily precipitation from Davis, California from March 2021 until January 2022 are shown. Data from: “DAVIS 2 WSW EXPERIMENTAL FARM, CA US USC00042294" www.ncei.noaa.gov.
